# Diatom Modulation of Microbial Consortia Through Use of Two Unique Secondary Metabolites

**DOI:** 10.1101/2020.06.11.144840

**Authors:** Ahmed A. Shibl, Ashley Isaac, Michael A. Ochsenkühn, Anny Cárdenas, Cong Fei, Gregory Behringer, Marc Arnoux, Nizar Drou, Miraflor P. Santos, Kristin C. Gunsalus, Christian R. Voolstra, Shady A. Amin

**Affiliations:** Marine Microbial Ecology Lab, Biology Program, New York University Abu Dhabi, Abu Dhabi, United Arab Emirates; International Max Planck Research School of Marine Microbiology, University of Bremen, Bremen, Germany; Department of Biology, University of Konstanz, Konstanz, Germany; Red Sea Research Center, Biological and Environmental Sciences and Engineering Division (BESE), King Abdullah University of Science and Technology (KAUST), Thuwal, Saudi Arabia; College of Resources and Environmental Science, Nanjing Agriculture University, Nanjing, China; Center for Genomics and Systems Biology, New York University Abu Dhabi, Abu Dhabi, United Arab Emirates; Center for Genomics and Systems Biology, Department of Biology, New York University, New York, NY, USA

**Author notes:** To whom correspondence should be addressed: Shady A. Amin.

**Keywords:** phycospheres, microbiomes, signaling, diatom, phytoplankton–bacteria interactions, secondary metabolites

## Abstract

Unicellular eukaryotic phytoplankton, such as diatoms, rely on microbial communities for survival despite lacking specialized compartments to house microbiomes (e.g., animal gut). Microbial communities have been widely shown to benefit from diatom excretions that accumulate within the microenvironment surrounding phytoplankton cells, known as the phycosphere. However, mechanisms that enable diatoms and other unicellular eukaryotes to nurture specific microbiomes by fostering beneficial bacteria and repelling harmful ones are mostly unknown. We hypothesized that diatom exudates may attune microbial communities and employed an integrated multi-omics approach using the ubiquitous diatom *Asterionellopsis glacialis* to reveal how it modulates its naturally associated bacteria. We show that *A. glacialis* reprograms its transcriptional and metabolic profiles in response to bacteria to secrete a suite of central metabolites and two unusual secondary metabolites, rosmarinic acid and azelaic acid. While central metabolites are utilized by potential bacterial symbionts and opportunists alike, rosmarinic acid promotes attachment of beneficial bacteria to the diatom and simultaneously suppresses the attachment of opportunists. Similarly, azelaic acid enhances growth of beneficial bacteria, while simultaneously inhibiting growth of opportunistic ones. We further show that the bacterial response to azelaic acid is widespread in the world’s oceans and taxonomically restricted to a handful of bacterial genera. Our results demonstrate the innate ability of an important unicellular eukaryotic group to modulate their microbial consortia, similar to higher eukaryotes, using unique secondary metabolites that regulate bacterial growth and behavior inversely in different bacterial populations.

## Introduction

Large swaths of eukaryotic lineages possess associated microbiomes that play central roles in maintaining host survival and ecological success (1). Several biotic and abiotic factors have been shown to drive microbiome assembly and modulation in special compartments and organelles of multicellular eukaryotes such as squid light organs (2), coral skeletons (3), mammalian guts (4), and roots and leaves of terrestrial plants (5). Contrarily, unicellular eukaryotes such as diatoms lack developmental features that can harbor microbes, yet rely heavily on essential bacterial growth factors (6–8) to proliferate and thrive in their environment. Diatoms are ubiquitous primary producers in aquatic environments that excrete up to 50% of their fixed carbon (9–11) into a diffusive boundary layer that surrounds individual cells. This physically sheltered microscale region, known as the phycosphere, is highly enriched in dissolved organic matter (DOM) and serves as the interface for diatom-bacteria associations (7, 12). Indeed, bacteria have been shown to heavily rely on phycosphere DOM to support their growth (13, 14) and must use motility, chemotaxis and/or attachment to chase and colonize the phycosphere (15). Recent research has shown that a variety of interactions spanning mutualism, commensalism, and parasitism occur between diatoms and specific groups of bacteria (7, 16–18). As single cells floating in aquatic environments, diatoms encounter beneficial (hereafter symbiotic) and opportunistic and algicidal (hereafter opportunistic) bacteria. However, the mechanisms that allow diatoms and other phytoplankton species to actively modulate incoming microbes to evade opportunistic bacteria and nurture symbiotic ones are mostly unknown. Due to the challenges of investigating phytoplankton–bacteria interactions in the field, most studies to date have relied on laboratory-controlled co-culture systems between phytoplankton and a single bacterium, an approach that has enriched our knowledge of phycosphere interactions but one that does not adequately mimic the microbial complexity in natural phycospheres. Here, we apply a holistic approach to a natural system derived from the environment by using multi-omics to show that DOM secretions by the globally widespread diatom *Asterionellopsis glacialis* (19) modulate microbial community behavior and growth. We hypothesize that diatom cells must adopt specific mechanisms to promote association with beneficial symbionts while repelling opportunists to offset the lack of specialized compartments to house microbiomes. To this end, *A. glacialis* strain A3 was cultivated from its natural environment, then freed of its associated bacteria and left to acclimate until the time of reseeding, marked by the re-introduction of its natural bacterial consortium to the diatom. Transcriptional and metabolomic changes in both the diatom and the bacterial consortium at different time points were assessed and potential representative symbiotic and opportunistic bacteria were cultivated from the consortium to further confirm hypotheses generated from multi-omics experiments.

## Results

To examine the interactions between the diatom and its bacterial consortium, we isolated *A. glacialis* A3 along with its natural microbial community (xenic *A. glacialis)* then cured it of bacteria using a suite of antibiotics to make it axenic, as described previously (20). After ~170 generations of acclimating the axenic *A. glacialis* A3 culture to the absence of bacteria, the true bacterial consortium composition was harvested by filtration from xenic cultures immediately before the reseeding experiment. At the time of reseeding, one portion of this natural bacterial community was added to the acclimated axenic *A. glacialis* A3 culture, generating a reseeded *A. glacialis* A3 treatment to investigate the response of the diatom to bacterial exposure and the response of bacteria to diatom exudates (Fig. S1). Two additional portions of the bacterial consortium were collected and used for shotgun metagenomics and metatranscriptomics (bacterial consortium control at 0.5 hours). Diatom transcriptomic samples (at 0.5 and 24 hours) were collected from the control axenic *A. glacialis* cultures and reseeded *A. glacialis* treatments. In addition, samples for metabolomics at two early (0.5 and 4 hours) and two late (24 and 48 hours) time points were collected (see Methods and Fig. S1).

The composition of the microbial consortium collected at the time of reseeding showed the dominance of six bacterial families, with Flavobacteriaceae comprising 38.9% of all metagenomic reads, followed by Rhodobacteraceae (16.6%), Erythrobacteraceae (16%), Alteromonadaceae (9.28%), Pseudomonadaceae (1.07%) and Oceanospirillaceae (1.03%) (Fig. 1*A*). To uncover how these families responded to diatom exudates, we assembled ten near-complete bacterial genomes from the microbial consortium metagenome. The metagenomically-assembled genomes (MAGs) belonged to most major families in the consortium, including Flavobacteriaceae (MAG9), Rhodobacteraceae (MAG3, MAG5, MAG6 and MAG11), Erythrobacteraceae (MAG10), Alteromonadaceae (MAG4 and MAG12), Oceanospirillaceae (MAG8), and Halomonadaceae (MAG13) (Table S1). Mapping metatranscriptome reads to all MAGs showed that the four Rhodobacteraceae MAGs recruited ~41% of mRNA reads and were responsible for the majority of differentially expressed genes (Table S2) at both early and late time points of reseeded samples relative to controls, despite representing ~10% of the bacterial consortium metagenome (Table S1).

**Figure 1.**
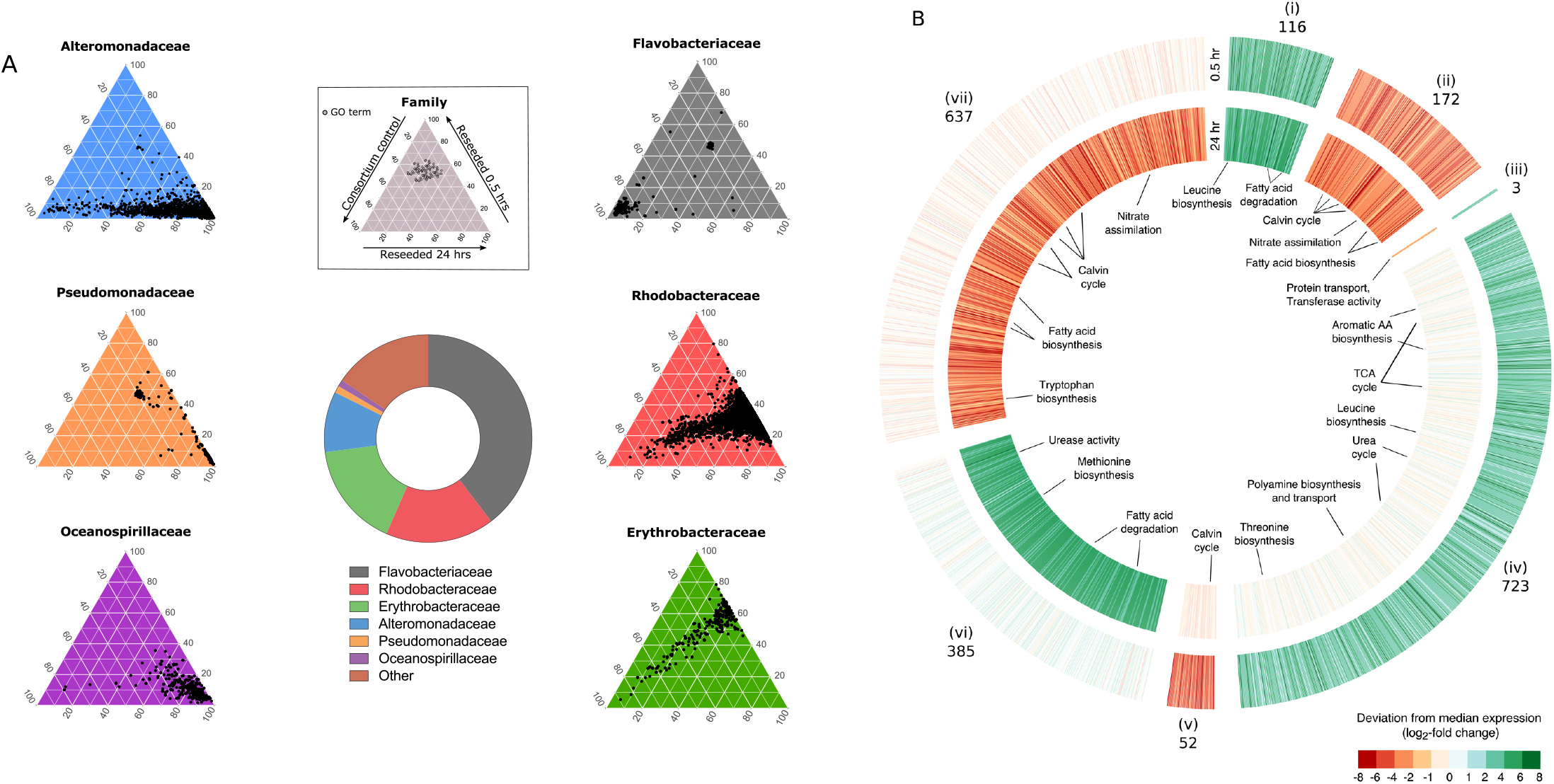
Major reprogramming of transcriptional responses of *A. glacialis* A3 and roseobacters in response to reseeding. (**A**) Central donut plot depicts relative abundances of the top six bacterial families in the consortium metagenomic dataset. Inset: key for color-coded ternary plots represent transcriptional responses of the bacterial families before (consortium 0.5 hours control) and after (reseeded 0.5 and 24 hours) reseeding based on biological triplicates. Each dot depicts a unique gene ontology (GO) annotation associated with transcripts from each of the six major families in the metatranscriptome. The position of each dot corresponds to the percent contribution of the sample (consortium control, reseeded 0.5 hours, and reseeded 24 hours) relative to the total normalized abundance of transcripts annotated with the same GO term, in copies per million (CPM). (**B**) Differentially expressed (DE) genes in reseeded *A. glacialis* A3 after 0.5 (outer circle) and 24 (inner circle) hours relative to axenic controls. Genes are organized into seven clusters (i-vii) based on their expression pattern at the two timepoints. Numbers indicate the number of DE genes in each cluster. Opaque clusters indicate genes that are not DE. TCA=tricarboxylic acid.

We examined the consortium metatranscriptome and confirmed that the Rhodobacteraceae (hereafter roseobacters) exhibited the most transcriptionally rapid and diverse response within 0.5 hours of reintroduction to the diatom, as evidenced by the large number of Gene Ontology terms associated with roseobacter genes expressed after reseeding. In stark contrast, other bacterial families either displayed no significant response to reseeding (Flavobacteriaceae), a decreasing response from 0.5 to 24 hours (Erythrobacteraceae), or were responsive only at 24 hours after reseeding (Alteromonadaceae, Pseudomonadaceae and Oceanospirillaceae) relative to the consortium control (Fig. 1 *A*).

The *A. glacialis* A3 transcriptome showed a major reprogramming of its transcriptional profile to differentially express ~14% of its protein-coding genes relative to axenic controls, coupled with temporal shifts in expression patterns (Fig. 1*B*). In response to consortium reseeding, transcripts for amino acid biosynthesis and fatty acid degradation were consistently upregulated, while nitrate assimilation, photosynthesis and carbon fixation were downregulated throughout the reseeding experiment. At 0.5 hours only, differentially upregulated *A. glacialis* A3 transcripts included those for spermidine biosynthesis and transport and the tricarboxylic acid (TCA) and urea cycles, while transcripts for methionine biosynthesis and urease activity were differentially upregulated at 24 hours only. We also observed differentially downregulated transcripts involved in the Calvin cycle at both 0.5 hours and 24 hours and tryptophan biosynthesis related transcripts downregulated at 24 hours only (Fig. 1*B* and Table S3).

The diatom and roseobacters transcriptional responses were coupled to major changes in the exometabolome. Exometabolomes sampled at two early and two late time points after reseeding (Fig. S1*B*) were analyzed using a quadrupole time-of-flight mass spectrometer (Dataset S1). The DOM landscape varied between axenic and reseeded samples (Fig. 2*A*). Interestingly, based on Mahalanobis distances (M_d_), the DOM composition at early time points was significantly more distinct from late time points in the reseeded samples (M_d_=3.88) than in axenic controls (Md=3.06) (Fig. *2B, C*), suggesting that DOM is temporally highly dynamic in response to consortium reseeding, similar to the diatom transcriptome. Analysis of the DOM elemental composition of extracted metabolites in axenic and reseeded samples using Fourier-transform ion cyclotron resonance mass spectrometry (FT-ICR-MS) (Datasets S2, S3) showed ~50% decrease in abundance of dissolved organic nitrogen (DON) in reseeded samples relative to axenic controls (Fig. S2).

**Figure 2.**
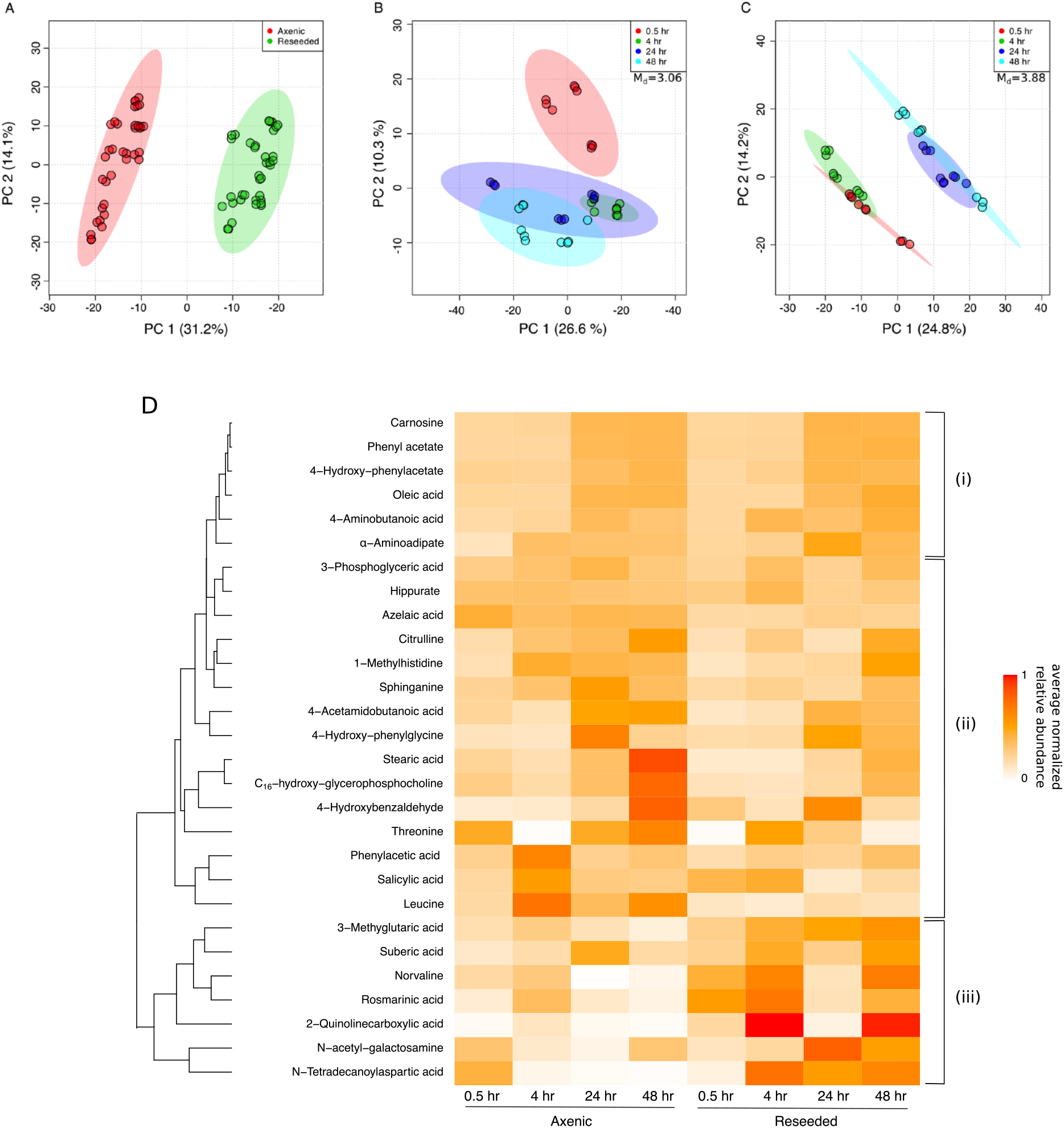
SPE-extracted DOM profile is highly influenced by reseeding. (**A-C**) Principal components analysis (PCA) plots of axenic and reseeded untargeted exometabolome samples. PCA was performed based on Mahalanobis distances (M_d_), comparing 1,237 SPE-extracted exometabolites between (**A**) axenic *vs.* reseeded samples, and (**B, C**) early (0.5 and 4 hours) and late (24 and 48 hours) timepoints for (**B**) axenic, and (**C**) reseeded conditions. Circles represent technical replicates (*n*=3) of three biological replicates. (**D**) Euclidean hierarchical clustering of 28 exometabolites (Table S4) identified in axenic and reseeded samples and confirmed using a library of in-house chemical standards. Colors represent average normalized relative abundance of each metabolite. (i) Prospective refractory diatom metabolites; (ii) Diatom metabolites possibly taken up by the consortium; (iii) Diatom metabolites with a potential signaling role.

The identity of 28 metabolites common in axenic and reseeded diatom samples was confirmed (Fig. 2*D* and Table S4) using an in-house chemical library of >660 molecules (see Methods), indicating these metabolites are secreted by the diatom. Most metabolites showed increasing relative abundance in axenic and reseeded samples as a function of time, but a markedly lower overall accumulation in reseeded samples relative to axenic controls (e.g., leucine, threonine, 3-phosphoglycerate), suggesting either diatom downregulation of the biosynthesis of these molecules in reseeded samples and/or bacterial uptake in reseeded samples. Bacterial uptake was corroborated by the transcriptional response of the diatom to reseeding, which showed upregulation of metabolite-specific biosynthesis genes and a concomitant upregulation of specific roseobacters transporters that take up these metabolites (Table S5). Seven metabolites showed significant increases in relative abundance in reseeded samples compared to axenic controls (e.g., rosmarinic acid), suggesting either a signaling role for these diatom metabolites or co-production by bacteria (Fig. 2*D*).

Based on the rapid response of the roseobacters to diatom exudates, we built a conceptual model of diatom-roseobacters interactions using the differential gene expression of *A. glacialis* A3, three roseobacters MAGs (MAG3, 5 and 6), and identified exometabolites (Fig. 3 and Tables S3, S4 and S5). In response to reseeding, the diatom upregulated genes involved in the biosynthesis of spermidine (log_2_-fc=3.8, *p*=0.09) and its transport (log_2_-fc=2.7, *p*=0.04) at 0.5 hours. Concomitantly, transcripts for spermidine uptake were overexpressed in both MAG3 (log_2_-fc=5.5, *p*=0.09) and MAG5 (log_2_-fc=6.2, *p*=0.08) at 0.5 hours. The diatom increased transcription of glutamate dehydrogenase at 0.5 (log_2_-fc=2.2, *p*=0.079) and 24 hours (log_2_-fc=5.4, *p*=0.006) to fuel the TCA cycle and/or the urea cycle, both of which were upregulated, by generating α-ketoglutarate and ammonia, respectively. Citrulline, a urea cycle intermediate released into the media, showed a differential decrease in abundance in reseeded samples versus axenic samples (*p*=0.007 at 24 hours; Fig. 2*D*), suggesting bacterial uptake. The diatom downregulated homologs of phosphoglycerate kinase (21) involved in the conversion of 3-phosphoglycerate (3-PGA) to glycerate 1,3-diphosphate in the plastid (log_2_-fc=-1.6, *p*=0.06) and cytoplasm (log_2_-fc=-5.3, *p*=0.06). 3-PGA transporters localized in the plastid were also downregulated at 0.5 hours after reseeding (log_2_-fc=-7.0 and −4.0, *p*=0.002 and 0.007, respectively), indicating no transport of 3-PGA across the plastid membrane and a buildup of 3-PGA in the cytoplasm. 3-PGA was released into the media and was presumably taken up by bacteria. Transporters for 3-PGA were not differentially expressed in MAG3, while a 3-PGA response regulator was overexpressed in MAG5 at 0.5 hours (log_2_-fc=5.3, *p*=0.097). Diatom transcripts involved in the biosynthesis of threonine were overexpressed at 0.5 hours (log_2_-fc=2.7, *p*=0.02) and transcripts involved in the biosynthesis of leucine were overexpressed at both 0.5 (log_2_-fc=5.2, *p*=0.001) and 24 hours (log_2_-fc=5.2, *p*=0.0003). Transporters likely involved in the extracellular secretion of both amino acids were either upregulated (threonine) at 0.5 hours (log_2_-fc=4.8, *p*=0.03) or not differentially expressed (leucine) at both time points. The secretion of threonine (*p*=0.03 and 0.05 at 0.5 and 4 hours, respectively) and leucine (*p*=0.003 at 4 hours) (Fig. 2*D*) into the media was concomitant with an upregulation of their transporters and subsequent assimilation of leucine into branched-chain fatty acid biosynthesis in the three roseobacters MAGs (Fig. 3 and Tables S3 and S5).

**Figure 3.**
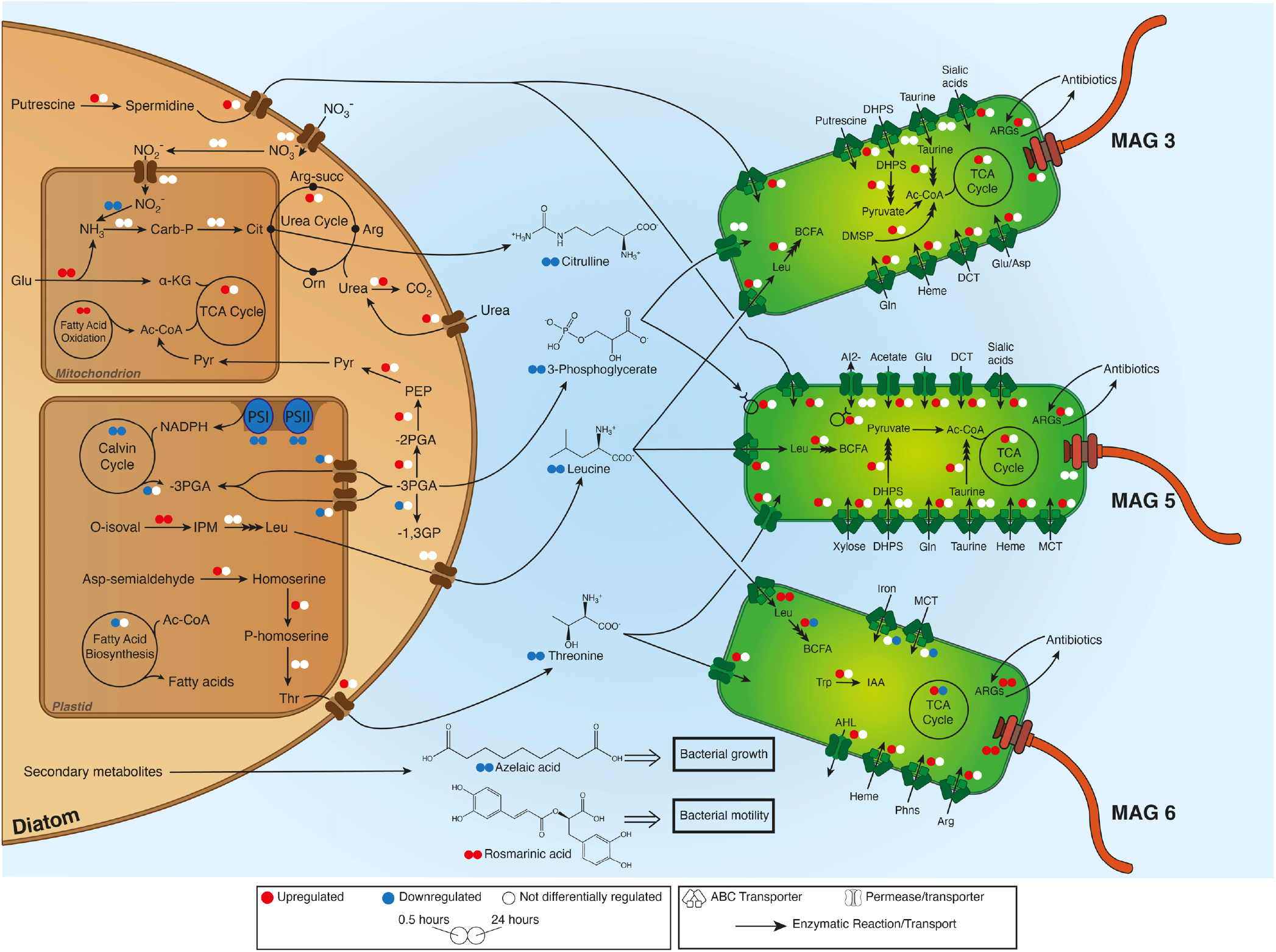
*A. glacialis* A3 preferentially promotes growth of roseobacters by secreting specific metabolites that influence bacterial growth and behavior. Summary of diatom-bacteria interactions highlighting the metabolic exchanges and differentially expressed (DE) genes in *A. glacialis* A3 and three roseobacters MAGs. Small colored circles (red: upregulation; blue: downregulation; white: no DE) represent differential expression of genes/processes at 0.5 (left) and 24 (right) hours after reseeding. Differential expression of metabolic cycles indicates that at least one gene was DE in one direction while no other genes were DE in the opposite direction. A complete list of genes and expression values are in Tables S3 and S5. Confirmed central and secondary molecules from the exometabolome (Table S4) are shown between the cells and their relative abundance is indicated by colored circles relative to axenic controls. Multiple stacked arrows indicate several enzymatic reactions. SAMamine=S-adenosylmethionineamine; Carb-P=carbamoylphosphate; Cit=citrulline; α-KG=α-ketoglutarate; Pyr=pyruvate; Arg-succ=argininosuccinate; Arg=arginine; Orn=ornithine; PEP=phosphoenolpyruvate; 3-PGA=3-phosphoglycerate; 1,3-GP=glycerate 1,3-diphosphate; PS=photosystem genes; O-isoval=o-isovalerate; IPM=isopropylmalate; Asp=aspartate; Glu=glutamate; Gln=glutamine; Leu=leucine; Thr=threonine; Trp=tryptophan; BCFA=branched-chain fatty acids; DHPS=2,3-dihydroxypropanesulfonate; ARGs=antibiotic resistance genes; DCT=dicarboxylate transporter; DMSP=dimethylsulfoniopropionate; AI-2=autoinducer-2; MCT=monocarboxylate 2-oxoacid transporter; Phns=phosphonates; AHL=acyl homoserine lactones; IAA=indole-3-acetate.

To confirm the ability of roseobacters to utilize diatom metabolites, we isolated bacteria from the bacterial consortium and sequenced their genomes (see Methods). Two isolates were identified as roseobacters species: *Sulfitobacter pseudonitzschiae* F5 and *Phaeobacter* sp. F10 and one isolate as an Alteromonadaceae species: *Alteromonas macleodii* F12. Phylogenomic analysis of isolate genomes and MAGs clustered *Phaeobacter* sp. F10 close to MAG6 (86.2% amino acid identity with *P. gallaeciensis*) (Fig. S3 and Table S6), while *A. macleodii* F12 clustered within the *A. macleodii* clade (Fig. S4 and Table S7). Subsequently, 16 diatom metabolites from Fig. 2*D* were used to test the ability of *S. pseudonitzschiae* F5 (a potential symbiont) and *A. macleodii* F12 (a potential opportunist) to utilize these metabolites as growth substrates. Despite the more rapid transcriptional responses of roseobacters to reseeding (Fig. 1*A*), both bacterial isolates were able to use most of these central metabolites as growth substrates (Fig. S5).

We sought to examine if diatom secondary metabolites can account for the advantage roseobacters have over other bacterial families in the microbial consortium, like the Alteromonadaceae. Cell attachment is an important mechanism used by bacteria to remain in the phycosphere to enhance access to diatom exudates (22). The motility of *S. pseudonitzschiae* F5, *Phaeobacter* sp. F10 and *A. macleodii* F12 was examined in the presence of a secondary metabolite not detected in diatoms before, rosmarinic acid, a common constituent of some terrestrial plants (23). Surprisingly, 2 μM rosmarinic acid significantly inhibited the motility of the symbionts *S. pseudonitzschiae* F5 and *Phaeobacter* sp. F10 and increased the motility of the opportunist *A. macleodii* F12 (Fig. 4). To confirm whether reduced motility enables the symbionts to attach to the diatom, *A. glacialis* A3 was co-cultured with each bacterial isolate. Indeed, *S. pseudonitzschiae* F5 and *Phaeobacter* sp. F10 exhibited strong attachment in the diatom phycosphere while *A. macleodii* F12 showed no apparent attachment (Fig. 4).

**Figure 4.**
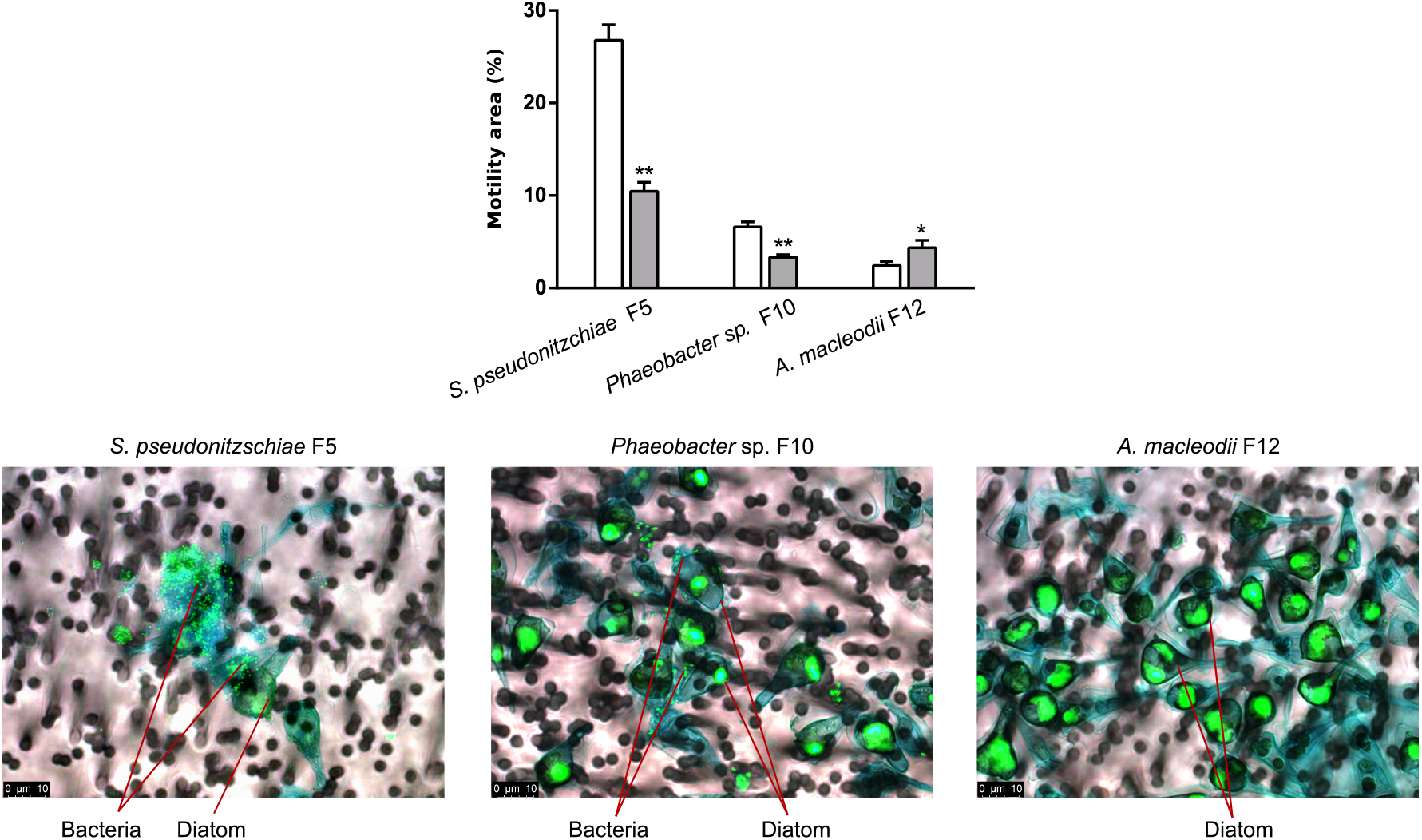
Diatom secondary metabolite rosmarinic acid reduces motility and promotes attachment of roseobacter symbionts to *A. glacialis* A3. Top: Motility behavior of strains *S. pseudonitzschiae* F5, *Phaeobacter* sp. F10, and *A. macleodii* F12 grown on semisolid (0.25% w/v) marine agar plates with (grey bars) or without (white bars) 2 μM rosmarinic acid. Error bars represent standard deviation (SD) of the three replicates. Significance was determined by Student’s *t*-test: \**p*<0.05 and \*\**p*<0.001. Bottom: Fluorescence microscopy images of co-cultures of the diatom with the two roseobacters strains and *A. macleodii* F12. SYBR Green I was used to visualize diatom and bacterial DNA; Alcian blue was used to stain the diatom exopolysaccharide matrix, known as transparent exopolymeric particles (TEP, in blue). Cocultures were gently filtered prior to microscopy onto 3-μm membrane filters to remove free-living bacteria. No bacteria are visible on TEP in the vicinity of diatom cells in the *A. macleodii* F12 panel, indicating that most *A. macleodii* F12 cells were free-living and were removed by gravity filtration.

In addition to rosmarinic acid, 100 μM azelaic acid, a byproduct of oleic acid metabolism, significantly inhibited the growth of *A. macleodii* F12 over a 24-hour period while the same concentration promoted growth of symbionts over a 48-hour period (Fig. 5*A-C*). Bacterial response to azelaic acid was shown to be controlled by a transcriptional regulator, AzeR (24). To shed light on the prevalence of the bacterial response to azelaic acid throughout the oceans, a Hidden Markov Model (HMM) profile of AzeR homologs detected in all three bacterial isolates was used to search the *Tara* Oceans database. AzeR homologs were consistently distributed at surface and deep chlorophyll maximum depths across the oceans, with most homologs belonging to Alteromonadales (19%) and Rhodobacterales (18%) (Fig. 5*D*). Mining the Pfam database for AzeR homologs indicated that the response to azelaic acid in publicly available bacterial genomes is mostly limited to the Proteobacteria phylum and is further restricted to six orders, including Alteromonadales and Rhodobacterales, to which the Alteromonadaceae and roseobacters belong, respectively (Fig. S6).

**Figure 5.**
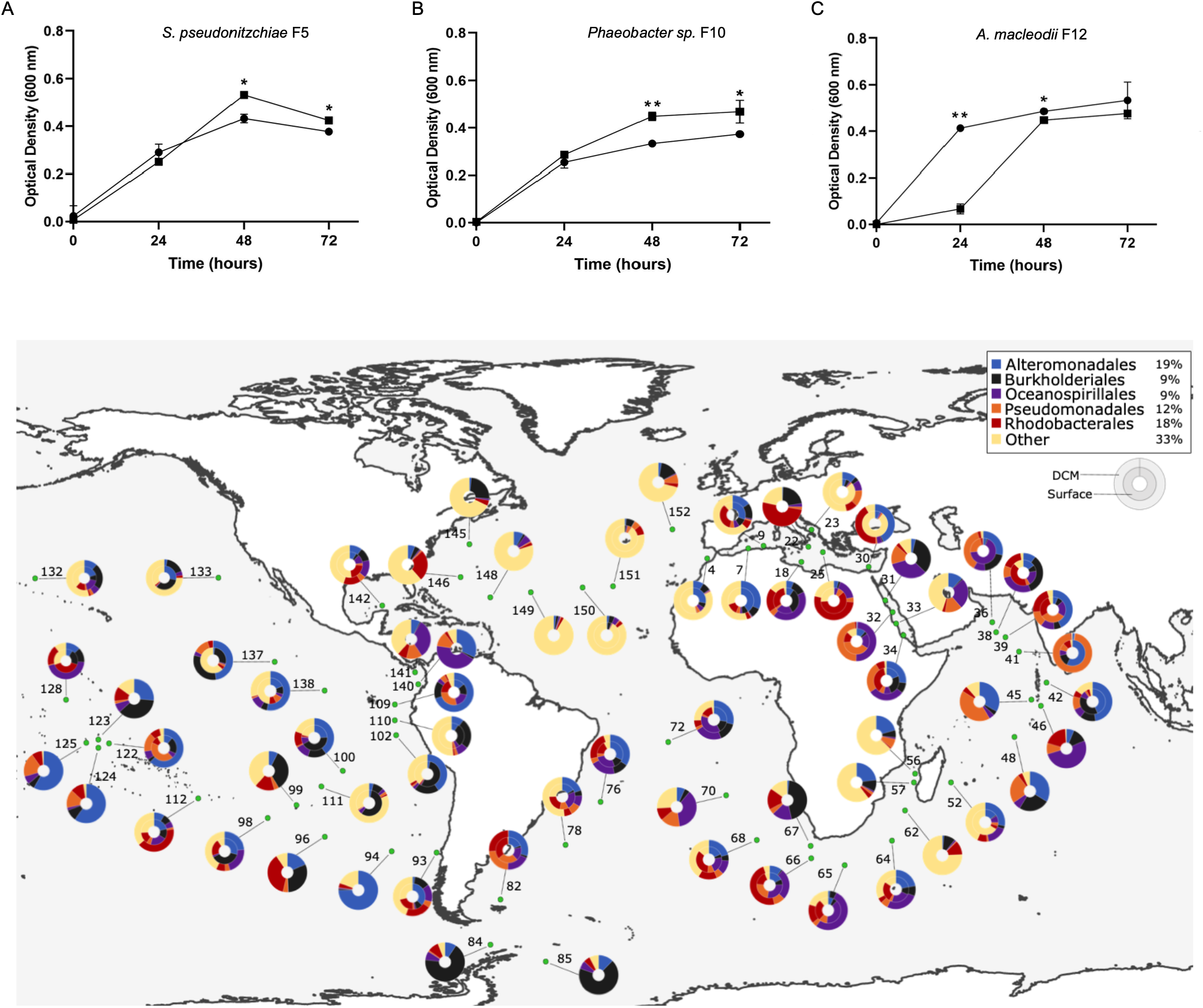
Diatom secondary metabolite azelaic acid promotes beneficial bacteria and controls potential opportunists. Growth of (**A**) *S. pseudonitzschiae* F5 and (**B**) *Phaeobacter* sp. F10, and (**C**) *A. macleodii* F12 on 10% marine broth supplemented with 100 μM azelaic acid (squares) compared to controls (circles). Error bars represent standard deviation (SD) of the three replicates. Significance was determined by Student’s *t*-test: \**p*<0.05 and \*\**p*<0.001. (**D**) Bacterial response to azelaic acid is geographically widespread throughout the oceans. The total percentage abundance of the azelaic acid transcriptional regulator, AzeR, homologs according to their taxonomic distribution is shown in the top right box. Rhizobiales makes up the majority of hits (39%) in the ‘Other’ group. The color-coded donut plots represent the percentage taxonomic abundance of AzeR homologs from all size fractions (0-3 μm) at the surface (inner circle) and deep chlorophyll maximum (outer circle) from the *Tara* Oceans Microbiome Reference Gene Catalog. Numbers refer to the *Tara* Oceans stations; single donut plots depict surface samples only.

## Discussion

Remineralization of phytoplankton-derived organic matter by heterotrophic bacteria plays a major role in the carbon cycle and accounts for the transformation of ~20 gigatons of carbon per year in the ocean’s euphotic zone (25). Our current understanding of the global passive and active release of DOM by phytoplankton has been largely studied in the context of primary production, grazing events, and virus-mediated cell lysis (26, 27). Still, underlying reasons for the active excretion by phytoplankton of significant amounts of low molecular weight organic compounds into the phycosphere (6) are still being debated (25, 28). Because of their microscopic size, transport of molecules around phytoplankton cells is mostly governed by diffusion, which leads to the accumulation of phytoplankton-derived DOM within the phycosphere (12). Bacteria in the ocean expend significant energy to track and colonize these DOM-rich hotspots to fuel their growth (29), employing a variety of mechanisms to succeed in the phycosphere, including establishing symbiotic exchanges with phytoplankton cells or producing algicidal agents that harm or kill phytoplankton (7, 30, 31). Therefore, it is imperative for phytoplankton cells to control the types of bacteria that come in contact with the phycosphere, as the outcome ultimately leads to survival or death. However, the mechanisms that enable ocean-drifting phytoplankton cells to attract beneficial bacteria and repel harmful ones in the phycosphere, if any, are mostly unknown.

The microbial community composition surrounding *A. glacialis* A3 is typical of bacteria associated with phytoplankton cultures and blooms (32, 33). Flavobacteria, the dominant lineage in the natural bacterial community associated with the diatom, often assimilate complex organic matter (e.g., polysaccharides) that require exoenzyme activity (34), especially during phytoplankton blooms (35), partially explaining their inactivity over shorter times with *A. glacialis* (<24 hours) (Fig. 1*A*). Within 0.5 hours of reintroducing the natural consortium to the axenic diatom culture, roseobacters rapidly dominated the bacterial transcriptional activity (Fig. 1*A*). The *Roseobacter* group spans >70 genera (36) with a highly versatile genetic repertoire (37, 38) that often dominate microbial assemblages surrounding particulate organic matter (39–41). They have been consistently shown to establish specific symbiotic relationships with diatoms (16, 17, 42) and are especially adept at acquiring phytoplankton-derived DOM (33, 43, 44). Despite their rapid response to *A. glacialis* A3 exudates relative to all other families, members of the *Roseobacter* group only represented 16.6% of the microbial consortium of the diatom, which is in line with the average *Roseobacter* group abundance in phytoplankton blooms (33). This discrepancy is potentially due to competition and chemical warfare between different bacterial taxa in the consortium, manifested by the overexpression of antibiotic resistance genes in all roseobacter MAGs (Fig. 3 and Table S5), which mitigates proliferation of any one bacterial group in the phycosphere. Indeed, production of diverse antimicrobial agents in a complex microbial community has been shown to maintain bacterial diversity (45), which explains why despite being the most active, roseobacters cannot solely dominate the phycosphere of *A. glacialis* A3. In contrast, Alteromonadaceae typically show strong algicidal activity against a wide range of phytoplankton lineages, including diatoms (46).

Isolation and sequencing of *S. pseudonitzschiae* F5, *Phaeobacter* sp. F10, and *A. macleodii* F12 from the natural diatom microbial consortium provides an ample prospect to better understand phytoplankton modulation of different bacterial taxa in the phycosphere. Remarkably, several *S. pseudonitzschiae* strains (16S rRNA sequence identity >97%) have been isolated from several diatom species originating from different oceanic regions (17, 47, 48). One such strain, *S. pseudonitzschiae* SA11, (clustered near *S. pseudonitzschiae* F5, Fig. S3) is a known diatom symbiont that enhances cell division of another diatom, *Pseudo-nitzschia multiseries*, via the hormone indole-3-acetic acid (17). Preliminary growth experiments between *S. pseudonitzschiae* F5 and *A. glacialis* A3 indicate that it also enhances *A. glacialis* cell division, similar to *S. pseudonitzschiae* SA11 and *P. multiseries* (Fei et al., *in review).* These findings suggest that *Sulfitobacter* is a conserved diatom symbiont. Interestingly, the close phylogenetic clustering of *Phaeobacter* sp. F10 and MAG6 indicate they are the same bacterium, whereas the placement of *A. macleodii* F12 within the group of the model bacterium *Alteromonas macleodii* (Fig. S4) suggests it is a common copiotrophic opportunist (49).

The conceptual model presented here (Fig. 3) clearly identifies the transcriptional and metabolomic responses of the host diatom and the surrounding roseobacters. The combination of multi-omics, bacterial isolation, and examination of the effects of different metabolites on these bacterial isolates provides several lines of evidence to support our conclusions. For example, upregulation of the biosynthesis of metabolites (Fig. 3) by the diatom in response to reseeding is corroborated by the detection of these metabolites in the exometabolome, bacterial transcriptional responses toward these metabolites supported by metatranscriptomics, and by growth experiments of bacterial isolates representing the *Roseobacter* group in the presence of these metabolites. Although we were not able to detect polyamines (e.g., spermidine) presumably produced by the diatom in our metabolome, the upregulation of genes involved in spermidine uptake by MAG3 and MAG5 suggests that these diatom N-rich molecules may be rapidly utilized by the roseobacters. Consistent with this observation, genes related to polyamine transformation were shown to be expressed mostly by roseobacters in coastal waters, where diatoms usually dominate phytoplankton composition (50). In addition to spermidine, the rapid depletion of DON relative to DOC in the reseeded exometabolome (Fig. S2) is supported by previous findings showing that labile N-containing compounds are preferentially utilized by roseobacters in estuarine waters (51). These observations suggest DON is more labile than dissolved organic carbon in the phycosphere. The significant decrease in abundance of another DON molecule, citrulline, after the reseeding of bacteria implies its potential uptake (Fig. 2*D*). Although citrulline has been shown to support bacterial growth as a sole carbon source, uptake mechanisms have not been yet identified (52), complicating our ability to confirm bacterial uptake. However, growth of *S. pseudonitzschiae* F5 on citrulline confirms the ability of members of the *Roseobacter* group to use it as a carbon source (Fig. S5).

Of 1,237 detected metabolites, we were able to confirm the presence of 28 using a custom-curated chemical library of >660 biomolecules (Fig. 2D). Many of these confirmed metabolites have never been shown to be produced by diatoms before, suggesting that diatoms may be a rich source of metabolites in the ocean. In addition to several central metabolites, we observe the release of obscure secondary metabolites such as quinolinecarboxylic acid, 3-methylglutaric acid, suberic acid, and carnosine (Fig. 2*D*), which may play a role in symbiotic interactions or defense with different marine bacteria. Interestingly, other confirmed metabolites that have not been shown to be produced by diatoms before, such as rosmarinic acid, azelaic acid, salicylic acid, hippurate, and N-acetyl-galactosamine (Fig. 2*D* and Table S4), are involved in plant defense and interkingdom signaling mechanisms (53, 54). Production and secretion of these metabolites by the diatom hints at a defense system response (55) akin to land plants. The significant shift in metabolic activity over time as the diatom host came in contact with the microbial consortia (Fig. 2*A-C* and Fig. S2) raises the question of the presence of more specialized compounds that potentially aid in shaping the phytoplankton microbiome. We sought to validate our hypothesis by examining the bacterial response to two of these secondary metabolites, rosmarinic acid and azelaic acid, using the isolated strains.

Rosmarinic acid was one of seven molecules that showed an increase in relative abundance within 0.5 hours of reseeding relative to axenic controls (*p*<0.05 at all timepoints) (Fig. 2*D*). This increase in abundance is due either to upregulation of its biosynthesis by the diatom in response to reseeding, suggesting an interkingdom signaling function, or due to bacterial co-production. Bacterial co-production can be ruled out given that rosmarinic acid is only known to be produced by some land plants and has never been shown to be produced by prokaryotes (56). We mined the diatom genome for rosmarinic acid biosynthesis genes using plant homologs but were unable to find any matches, suggesting that diatoms may use a unique biosynthesis pathway different from legumes. Interestingly, rosmarinic acid significantly suppressed motility and promoted attachment of symbionts but had the opposite effect on *A. macleodii* F12 (Fig. 4). Rosmarinic acid was recently reported to be produced by *Arabidopsis thaliana* as a mimic of pathogenic bacterial quorum sensing autoinducers (57). It is likely that rosmarinic acid is also interfering with bacterial quorum sensing to control bacterial motility and attachment in the phycosphere, a hypothesis that appears to be supported by recent findings (Fei et al., *in review).*

Azelaic acid, a C9-dicarboxylic acid and a byproduct of oleic acid metabolism, is also produced by the diatom (Fig. 2D). Azelaic acid primes plant defenses (58) and leads to the production of another defense signal, salicylic acid (59), which is also released by the diatom (Fig. 2*D*). The decrease in abundance of azelaic acid in reseeded exometabolomes (*p=*0.0002 and 0.003 at 4 and 48 hours, respectively; Fig. 2*D*) and its influence on growth of bacterial isolates suggest that the compound was assimilated by roseobacters and Alteromonadaceae. A congener of azelaic acid, suberic acid (C8-dicarboxylic acid), is also produced by the diatom and promotes the growth of *S. pseudonitzschiae* F5 and *A. macleodii* F12 alike (Fig. S5). The similar structure and activity of both congeners suggest that azelaic acid targets growth of Alteromonadaceae while suberic acid may target other bacteria by inhibiting their growth. While such a strategy may enable diatoms to modulate different bacterial groups, roseobacters gain an apparent advantage by utilizing a wide range of substrates from diatoms. Analysis of transporters in the genomes *of S. pseudonitzschiae* F5*, Phaeobacter* sp. F10 *and A. macleodii* F12 indicate that the roseobacters possess a significantly higher number of transporters normalized to genome size relative to *A. macleodii* F12 (Fei et al. in review). The mechanism of growth inhibition and promotion by azelaic acid remains unknown and further work is needed to reveal its mechanism of action.

Recent findings show that bacterial community assembly in synthetic phycospheres can be predicted from the linear combination of taxa supported by growth on single phytoplankton central metabolites (60). Our findings further expand on our understanding of the role of metabolites in the phycosphere by incorporating host response to presence of different bacterial groups, manifested in the secretion of two unique secondary metabolites. Secretion of secondary metabolites by multicellular eukaryotes to modulate their microbiomes has been broadly reported (61–63). The ability of diatoms (and presumably other unicellular eukaryotes) to exert control over their microbial associates, indicating a capacity to nurture microbiomes, may have evolved earlier than the rise of multicellularity in eukaryotes. More interestingly, the ability of diatom-derived metabolites to have opposite phenotypic and/or behavioral effects on two different bacterial populations, to our knowledge, has not been widely shown. This ability hints at complex evolutionary trajectories of how diatoms evolved the use of these metabolites and the role of secondary metabolism in interkingdom signaling. Further work is needed to characterize the mechanisms of action of these unique molecules in bacteria and to further identify other diatom metabolites and their role in modulating bacterial populations. Shedding light on these mechanisms has the potential to expand our understanding of food web dynamics and the role of phycosphere bacteria in carbon cycling.

## Summary

Multicellular eukaryotes use diverse strategies to recruit and modulate microbiomes in specialized developmental organelles, such as the mammalian gut (64). In contrast, unicellular eukaryotes such as diatoms lack specialized organelles to house microbiomes, and despite numerous observations that they possess unique microbial communities (65–67), it is not clear how they can modulate transient microbes. We show that in addition to phytoplankton-derived central metabolites accessible to bacteria, the diatom *A. glacialis* A3 employs unique secondary metabolites to promote the proliferation of beneficial bacteria and demote opportunists. The functional roles of signaling of secondary metabolites in marine environments are an important piece of the puzzle linking symbiotic exchanges between phytoplankton and bacteria with carbon cycling in the euphotic zone. Although signaling molecules are believed to constitute a minor fraction of DOM in the euphotic zone, their regulation of microbial metabolism and growth means they can exert a major influence on carbon cycling. This study provides a glimpse into the potential evolution of molecules from the same algal source that have opposite effects on two different groups of bacteria but a favorable outcome for the host. Such an efficient strategy to achieve two outcomes on symbionts and non-symbionts in the diverse euphotic zone (6, 68) hints that microalgae and other unicellular eukaryotes modulate microbiomes.

## Methods

### Diatom isolation and growth

*Asterionellopsis glacialis* strain A3 was isolated from the Persian Gulf and identified as described previously (20). All cultures were maintained in *f/2+Si* medium (69) in semi-continuous batch cultures (70) and incubated in growth chambers (Percival, Perry, IA) at 22°C, 125 μE m^-2^ s^-1^, and a 12:12 light/dark cycle. Light flux was measured using a QSL-2100 PAR Sensor (Biospherical Instruments Inc., San Diego, CA). Growth was monitored by measuring *in vivo* fluorescence using a 10-AU fluorometer (Turner Designs, San Jose, CA). All cultures were acclimated throughout the experiments as described below for at least three transfers using semi-continuous batch cultures. Cultures were considered acclimated if the growth rates of three consecutive transfers of triplicate cultures did not vary by more than 15%. Specific growth rates *(μ)* were calculated from the linear regression of the natural log of *in vivo* fluorescence versus time during the exponential growth phase of cultures. Standard deviation of *μ* was calculated using *μ* values from biological replicates over the exponential growth period.

### Microbial consortium reseeding experimental design

To examine the interactions between *A. glacialis* A3 and its bacterial consortium, the diatom was first made axenic as described previously (20). In brief, approximately 25 mL of a late-exponential phase growing *A. glacialis* A3 culture was gravity filtered onto a 0.65-μm pore-size polycarbonate membrane filter (Millipore). Cells were quickly rinsed with sterile *f/2+Si* media. Using sterile tweezers, the filter was removed from the filtration unit and washed for ~1 min in sterile media containing 20 mg/mL Triton X-100 detergent to remove surface-attached bacteria. The filter was discarded after re-suspension of cells by gentle shaking in sterile detergent-free media. Cells were again gravity filtered onto a fresh 0.65-μm pore-size polycarbonate membrane filter and rinsed with sterile media. Subsequently, cells were washed off the filter by gentle shaking into sterile media containing a suite of antibiotics (per mL: 50 μg streptomycin, 66.6 μg gentamycin, 20 μg ciprofloxacin, 2.2 μg chloramphenicol, and 100 μg ampicillin). Cells were then incubated in antibiotic-containing media for 48 hours under regular growth conditions. Finally, 0.5–1.0 mL of antibiotics-treated cells was transferred to antibiotic-free media. Cultures were regularly monitored for bacterial contamination by checking for bacterial growth in Zobell marine broth 2216 (HiMedia) (71) in addition to filtering 2-3 mL of exponential-phase growing culture and using Sybr Green I (Invitrogen) staining and epifluorescence microscopy (Nikon Eclipse 80i) as described previously (72). This axenic *A. glacialis* A3 culture was left to acclimate to no bacteria for ~170 generations and was subsequently used for the reseeding experiment.

To conduct the reseeding experiment, axenic and xenic *A. glacialis* A3 cultures were acclimated to growth in 1 L batch cultures. Xenic and axenic *A. glacialis* A3 cultures were grown side by side to allow the harvesting of the true bacterial consortium composition and adding it to the axenic *A. glacialis* A3. To begin the experiment, 6 L of axenic and 9 L of xenic *A. glacialis* A3 culture batches were inoculated at the same cell density (~ 5,000 diatom cells/mL) and time. Once both cultures reached a diatom cell density of ~1×10^5^ cells/mL, the xenic cultures were pooled, gently sonicated to detach diatom-attached bacteria and filtered through a sterile 3-μm polycarbonate filter (25 mm, Whatman, NJ, United States) to remove diatom cells; the filtrate containing the microbial community was collected in a sterile flask and was used for subsequent steps. The filtrate was subsequently centrifuged at 4,000 rpm for 20 minutes using an Avanti J-26 XPI centrifuge (Beckman Coulter, Inc.) to concentrate the bacterial consortium and remove residual organic carbon from the media. The bacterial pellet was washed with sterile *f/2*+Si once, centrifuged and subsequently reconstituted in 3 mL of sterile media. Bacterial cell density was enumerated using epifluorescence microscopy (Nikon Eclipse 80i) as described previously (72). This bacterial consortium stock was divided into three parts, each containing ~ 9 × 10^5^ cells/mL; 1) a third of the sample was used to isolate DNA for bacterial consortium metagenomics; 2) a third of the sample was incubated in triplicate 250 mL sterile *f/2+Si* media for 0.5 hours [this sample was used for the bacterial consortium RNA control with no diatom (bacterial consortium control)]; 3) the remainder of the sample was added to triplicate 1 L bottles of the acclimated axenic *A. glacialis* A3 (reseeded diatom). The remainder of the axenic *A. glacialis* A3 culture (3 L) served as triplicate axenic control (axenic diatom). This scheme ensured that the reseeded diatom samples contained similar diversity and density of bacteria and diatom relative to the original xenic culture. We avoided adding bacteria from natural seawater to ensure our experiments included originally isolated diatom symbionts. The beginning of the reseeding experiment (*t*=0) is marked by the addition of the bacterial consortium to the axenic diatom. A simplified schematic of the experimental design and diatom-bacterial consortium growth is shown in Fig. S1A.

### Diatom DNA and RNA isolation and sequencing

DNA was isolated from axenic *A. glacialis* A3 by filtering cells onto a 3-μm polycarbonate filter and using the Wizard SV Genomic DNA Purification kit (Promega) following the manufacturer’s instructions and quantified on a Qubit 3.0 fluorometer using the DNA high sensitivity assay kit (Thermo-Fisher Scientific). The diatom DNA library was prepared with 200 ng of starting material using the TruSeq DNA Nano kit (Illumina, San Diego, CA, USA).

For axenic diatom RNA samples, cultures were filtered through 3-μm polycarbonate filters (25 mm, Whatman, NJ, United States) at 0.5 hours and 24 hours after the beginning of the reseeding experiment. For reseeded diatom samples, cultures were filtered through 3-μm polycarbonate filters to obtain diatom-enriched samples then through 0.2-μm polycarbonate filters to obtain bacterial consortium samples (Fig. S1A). All filters were flash frozen in liquid nitrogen and later stored in −80°C until further processing. From the 3-μm filters, cells were lysed by bead beating with sterile beads (Sigma) for 10 minutes followed by total RNA isolation using the ToTALLY RNA total RNA Isolation kit (Ambion) according to the manufacturer’s instructions. Samples were treated with two rounds of DNase to remove contaminating DNA using Turbo-DNase (Ambion). Ribosomal RNA (rRNA) were removed using the Poly(A)Purist MAG kit (Thermo-Fisher Scientific) following the manufacturer’s instructions. The mRNA was amplified using a MessageAmp II aRNA Amplification kit (Invitrogen) according to the manufacturer’s instructions. RNA libraries were prepared with a maximum of 50 μL starting material, as per protocol instructions, using the TruSeq RNA v2 kit (Illumina, San Diego, CA, USA).

The resulting libraries’ concentrations and size distributions were assessed on a Qubit 3.0 fluorometer using the DNA high sensitivity assay kit (Thermo-Fisher Scientific) and a Bioanalyzer 2100 (Agilent, Santa Clara, CA, USA). Following this, libraries were normalized, pooled and quantified by qPCR with the KAPA Library quantification kit for Illumina platforms (Kapa Biosystems, Wilmington MA, USA) on a StepOnePlus qPCR system (Thermo-Fisher Scientific). One replicate library, B7, belonging to the reseeded bacterial samples at 24 hours was discarded due to low quality. Finally, samples were loaded at 12 pM with 2% phiX on a High Output FlowCell and paired-end sequenced (2×100 bp) on the Illumina HiSeq 2500 platform available at the NYU Abu Dhabi Center for Genomics and Systems Biology (Table S8).

### Diatom genome

Raw genomic reads were assessed with the FastQC v0.11.5(73) tool. Low-quality bases and sequencing adaptor contaminants were removed by the Trimmomatic v0.36 tool (74) with the following parameters: “ILLUMINACLIP:adapter.fa:2:30:10 TRAILING:3 LEADING:3 SLIDINGWINDOW:4:15 MINLEN:36”. Quality trimmed reads were then *de novo* assembled on Platanus v1.2.4 (75) to yield 1,840 scaffolds of size >10kb out of 6,925 scaffolds with N50=21,686 and a total size of 66.5 Mbp. The final assembly was assessed for accuracy and completeness with QUAST v5.0.2 (76), and BUSCO v3 (77).

### Diatom transcriptome

Raw RNAseq reads were quality trimmed as described for the genome. HISAT2 v2.0.4 (78) was used to map the reads to the assembled genome. Generated SAM files were converted to alignment files in BAM format and sorted by coordinates with SAMtools v1.5 (79). Using StringTie v1.3.0 (80), GTF files per sample were created then merged into one file representing the transcriptome. Transcript assemblies were annotated on the Trinotate pipeline (http://trinotate.github.io) following Bryant *et al* (81). Significant differences in gene expression between the samples were evaluated with DESeq2 v1.14.1 (82) at a false discovery rate (FDR) of 0.1 and a minimum log_2_-fold change of 0.5. Subcellular localization of gene products was determined on DeepLoc (83). Differentially expressed genes across different timepoints were visualized using the Circos package (84).

### Bacterial consortium DNA and RNA isolation and sequencing

For bacterial consortium metagenomic samples, bacterial DNA was isolated from the bacterial consortium control sample using bead-beating with sterile beads (Sigma) for 10 minutes followed by the EZNA Bacterial DNA kit (Omega Bio-Tek) according to the manufacturer’s instructions and then quantified on the Qubit 3.0 fluorometer using the DNA high sensitivity assay kit (Thermo-Fisher Scientific). The consortium metagenomic library was prepared with 200 ng of starting material using the TruSeq DNA Nano kit (Illumina, San Diego, CA, USA) and paired-end sequenced (2×100 bp) on the Illumina HiSeq 2500 platform.

For bacterial consortium RNA samples, consortium control samples were filtered through 0.2-μm polycarbonate filters (25 mm, Whatman, NJ, United States) at 0.5 hours after the beginning of the reseeding experiment. Reseeded *A. glacialis* A3 cultures were filtered through 3-μm then 0.2-μm polycarbonate filters at 0.5 hours and 24 hours after the beginning of incubation (Figure S1B). All filters were flash frozen in liquid nitrogen and later stored in −80°C until further processing. From the 0.2-μm filters, cells were lysed by bead beating with sterile beads (Sigma) for 10 minutes followed by total RNA isolation using the RNeasy Mini kit (Qiagen, Germantown, MD) according to the manufacturer’s instructions. Samples were treated with two rounds of DNase to remove contaminating DNA using Turbo-DNase (Ambion). Ribosomal RNA (rRNA) were removed using the MicrobExpress Bacterial mRNA enrichment kit (Ambion) following the manufacturer’s instructions. mRNA was amplified using the MessageAmp II-Bacteria RNA Amplification kit (Invitrogen) according to the manufacturer’s instructions. RNA libraries were prepared with a maximum of 50 μL starting material, as per protocol instructions, using the TruSeq RNA v2 kit (Illumina, San Diego, CA, USA). The resulting library sizes and distributions were assessed on the Bioanalyzer 2100 (Agilent, Santa Clara, CA, USA). Following this, libraries were normalized, pooled and quantified by qPCR with the KAPA Library quantification kit (Illumina, San Diego, CA, USA). Finally, samples were loaded at 12 pM with 2% phiX on a High Output FlowCell and paired-end sequenced (2×100 bp) on the Illumina HiSeq 2500 platform (Table S8).

### Bacterial consortium metagenome, binning and assembly

Raw metagenomic reads were quality trimmed on Trimmomatic v0.36, with a minimum length of 75 bp. Quality-checked reads were mapped to the *A. glacialis* A3 genome on BBtools using the BBmap package v37.10 (http://sourceforge.net/projects/bbmap/) with default parameters. Reads that did not map to the diatom were then used as input for Kaiju v1.5.0 (85) to determine the taxonomic profile and abundance of the microbial community at the protein level. De novo assembly was done using MEGAHIT v1.0.2 (86) with a k-*mer* size of 127 and scaffolds were binned into metagenomically-assembled genomes (MAGs) on MetaBAT v0.25.4 (87). The MAGs were assessed for completeness and contamination with CheckM v1.0.7 (88) then visualized and refined on Anvi’o v3 (89) until contamination values dropped below 5%. The closest genomic neighbor was determined by performing whole-genome comparisons of amino acid identities (AAI) using the Microbial Genomes Atlas (MiGA) (90) (Table S1). Functional annotation of the MAGs was performed on Prokka v1.12 (91).

### Bacterial consortium metatranscriptomes

Raw RNAseq reads were quality trimmed as described above. Paired-end reads were merged on Flash v1.2.11 (92) and rRNA fragments were identified and removed using SortMeRNA v2.0 (93). Non-rRNA reads were mapped to protein-coding genes of the MAGs with Bowtie2 v2.3.3 (94) (Table S2). Resulting SAM files were used to quantify gene expression levels using eXpress v1.5.1 (95) considering only genes with a minimum read count of 10 per group. Significant differences in gene expression between the samples were evaluated with DESeq2 v1.14.1 at a false discovery rate (FDR) of 0.1 and a minimum log_2_-fold change of 0.5. To infer the functional potential of the entire bacterial consortium, functional profiling against UniRef50 (96) was performed using HUMAnN2 v0.11.2 (97). Gene families were further mapped to Gene Ontology (GO) terms (98) and structured into pathways with MetaCyc (99) to generate “copies per million (CPM)” values across the different conditions. Data plots were generated in R v3.4.3 (100) with RStudio v1.2.1335 (RStudio Inc., Boston, MA, USA) and packages ggplot2 v3.1.1 (101) and ggtern v3.1.0 (102).

### Exometabolite extraction

All glassware used was acid washed (1.2 M HCl), rinsed with MilliQ-H_2_O, furnace-baked at 420°C and sterilized for a minimum of 12 hours to eliminate residual organic carbon contamination. All solutions were made with either MilliQ-H_2_O or LC-MS grade methanol (Thermo-Fisher). Cell-free filtrates from the axenic diatom and reseeded diatom samples at all timepoints (i.e. 0.5, 4, 24 and 48 hours, figure S1) were placed in 500-mL dark glass bottles (Thermo-Fisher), acidified to pH ~3 using 100% formic acid (Sigma). No bacterial consortium control was used because the consortium stock culture was free of carbon and would not survive. For QToF-MS, organic molecules were extracted by passing each replicate onto 500 mg Oasis HLB solid-phase extraction (SPE) cartridges (Waters, USA) using a peristaltic pump (MasterFlex Easy-Load 3, USA) at a flowrate of ~5 mL/min. All SPE cartridges were pre-conditioned according to the manufacturer’s instructions. Salts were washed from the SPE cartridges using 0.1% trifluoroacetic acid in MilliQ-H_2_O. Organic molecules were eluted into 5-mL borosilicate tubes (Thermo-Fisher) using 5% ammonium hydroxide in methanol. For FT-ICR-MS, organic molecules were extracted by passing samples as described for Q-ToF-MS except for the use of PPL Bond-Elut solid-phase extraction columns (Agilent Technologies, US), according to the manufacturer’s instructions, instead of Oasis HLB. All extracts were immediately dried using a Savant SC210A SpeedVac concentrator (Thermo-Fisher) and stored at −80°C until analysis.

### UHPLC-QToF-MS

Metabolites were analyzed on a Bruker Impact II HD quadrupole time-of-flight mass spectrometer (QToF-MS, BrukerDaltonik GmbH, Bremen, Germany) coupled to an Agilent 1290 UHPLC system (Agilent, US). Metabolites were separated using a reversed-phase (RP) method, where medium-polarity and non-polar metabolites were separated using an Eclipse Plus C18 column (50mm × 2.1mm ID) (Agilent, US). Chromatographic mobile phases consisted of MilliQ-H_2_O + 0.2% formic acid (buffer A), Acetonitrile + 0.2% formic acid (buffer B). The gradient started with 95% A and 5% B, with a gradient of 18 min to 100% B and 2 min at 100% B. Every run was followed by a 5-min wash step from buffer B to buffer A to isopropanol and back to the initial condition, where the column was equilibrated for another 2 mins. Detection was carried out in positive and negative ionization modes with the following parameters: ESI settings: dry gas temperature = 220 °C, dry gas flow = 8.0 L/min, nebulizer pressure = 2.2 bar, capillary voltage = 4500 V, end plate offset = (-)500 V; MS-ToF setting: Funnel 1 RF = 150 Vpp, Funnel 2 RF = 200 Vpp, Hexapole RF = 50, Quadrupole Ion Energy = 1 eV, Collision Energy = 7 eV, untargeted MS/MS = stepping 30 - 50 eV; Acquisition Setting: mass range = 50 - 1300 m/z, Spectra rate = 6.0 Hz spectra/s, 1000 ms/spectrum. Auto MS/MS was performed in stepping mode, splitting each fragmentation scan equally into 25 and 50 eV.

Calibration, retention time alignment and peak picking of individual LC-MS runs were performed using the T-Rex 3D algorithm of Metaboscape v4.0 (BrukerDaltonik GmbH, Bremen, Germany). Background noise was removed by applying an intensity threshold of 1000. Peak-picking and integration were accompanied by ^13^C cluster detection to verify molecular features and remove those which appear in less than 40% of samples. Peak annotation was performed using an in-house generated spectral library of 668 biomolecules from the Mass Spectrometry Metabolite Library of Standards (IROA Technologies, US) and further using the Bruker Personal MS/MS Library (BrukerDaltonik GmbH, Bremen, Germany). The acquired LC-MS data was normalized according to sample volume, scaled across all samples and log10-transformed. Multivariate statistical analysis was performed using MetaboAnalyst v3.0 (103) on >1,200 metabolites (Dataset S1), including confirmed metabolites, to generate principal component analysis (PCA) plots for axenic vs. reseeded conditions at all timepoints. Mahalanobis distances were calculated on R v3.4.3. The heatmap of confirmed metabolites (Table S3) was visualized using the ComplexHeatmap package (104). Significance in relative abundance between different time points in reseeded and axenic samples were calculated using a Student’s t-test (p<0.05).

### FT-ICR-MS

Fourier-transform ion cyclotron resonance mass spectrometry (FT-ICR-MS) was used to determine the molecular composition of dissolved organic matter (DOM) components in the exometabolome. High-resolution mass spectra were acquired on a SolariX FT-ICR-MS (BrukerDaltonik GmbH, Bremen, Germany) equipped with a 7 Tesla superconducting magnet and Paracell analyzer. Samples were directly injected into an electrospray ionization (ESI) source (BrukerDaltonik GmbH, Bremen, Germany) at a flow rate of 2 μL/min operated in negative ionization mode with a capillary voltage = 4500 V, end plate offset = (-)500 V, nebulizer pressure = 2 bar, dry gas flow = 10 L/min, dry gas temperature = 220°C. Spectra were acquired with a time domain of four mega words in 2ω resonance mode over a mass range of m/z 80 to 1000, with an optimal mass range from 200-600 m/z. Three-hundred scans were accumulated for each sample. Spectra were internally calibrated with a fatty acids reference list on the DataAnalysis 5.0 software (Bruker, Germany). Peak alignment was performed with maximum error thresholds of 0.01 ppm. The FT-ICR-MS spectra were exported to peak lists with a cut-off signal:noise ratio of 3 and a minimal signal intensity of 10^6^. Chemical formulae calculation was performed with an error threshold of 0.5 ppm from the exact mass for the chemical formula and isotopic fine structure. Chemical formulae were only generated if all theoretical isotope peaks (100%) were found in spectra (Datasets S2 and S3).

### Bacterial isolation, genomic DNA extraction, sequencing and assembly

To isolate individual bacterial strains from the bacterial consortium, 200 μL of the xenic *A. glacialis* A3 culture in log-phase diluted in sterile seawater were spread evenly on Zobell marine agar 2216 (HiMedia) plates and incubated at 25°C in the dark. For further purification, single colonies were picked, restreaked onto new agar plates and incubated as before. Cells were subsequently inoculated into marine broth and incubated at 28°C in a shaker incubator at 180 rpm. 2 mL of three bacterial cultures at an OD_600_ of 1 were centrifuged at 5000 rpm for 10 minutes to pellet the cells. Genomic DNA was extracted with the EZNA Bacterial DNA kit (Omega Bio-Tek) following the manufacturer’s instructions and quantified on the Qubit 3.0 fluorometer using the DNA high sensitivity assay kit (Thermo-Fisher Scientific). Genomes of the bacterial isolates were sequenced using Illumina MiSeq and PacBio platforms at either Apical Scientific (Selangor, Malaysia) or Novogene Bioinformatics Technology Co., Ltd (Beijing, China). PacBio reads were assembled into contigs using Canu v1.7 (105) after trimming and filtering. Raw reads were further mapped to the primary assemblies to identify and correct errors with BLASR v5.3 (106) and Arrow v2.2.1 (SMRT Link v7.0, www.pacb.com). Illumina paired-end 150 bp reads were trimmed using BBDuk and aligned with BBmap (https://sourceforge.net/projects/bbmap/) for further polishing of the PacBio assemblies. Resulting datasets were used as input to Pilon (107) for error correction and genome assembly improvement. The final consensus reference genomes *Sulfitobacter pseudonitzschiae* F5, *Phaeobacter* sp. F10 (Rhodobacteraceae) and *Alteromonas macleodii* F12 (Alteromonadaceae) were annotated on Prokka v1.12 and checked for completeness using BUSCO v3.

### Phylogenomics

To investigate the phylogenetic placement for roseobacter MAGs and the two isolated strains *S. pseudonitzschiae* F5 and *Phaeobacter* sp. F10, 43 complete genomes from the Rhodobacteraceae family and *Agrobacterium tumefaciens* Ach5, used as an outgroup, were downloaded from NCBI (Table S6). To investigate the phylogenetic placement for Alteromonadaceae MAGs and the isolated strain *A. macleodii* F12, 20 complete genomes from the Alteromonadaceae family and *Pseudomonas syringae* CC1557, used as an outgroup, were downloaded from NCBI (Table S7). First, bcgTree (108) was used to concatenate sequences of 107 single-copy core genes, located by HMMER v3.1b2 (109). MUSCLE v3.8.31 (110) and Gblocks 0.91b (111) were used to create and refine a multiple sequence alignment, respectively. The ETE3 package (112) was implemented on the final alignment using RAxML (113) with a JTT+GAMMA substitution model and 1000 bootstraps to generate the phylogenomic trees.

### Bacterial growth assays

To assess the effects of diatom metabolites on the growth of bacteria, a representative of Rhodobacteraceae strains (*S. pseudonitzschiae* F5) and *A. macleodii* F12 were tested for growth on citrulline, norvaline, azelaic acid, leucine, threonine, hippurate, carnosine, 3-quinolinecarboxylic acid, salicylic acid, suberic acid, phenylacetic acid, 4-hydroxybenzaldehyde, 1-methylhistidine, 3-phosphoglyceric acid and phenyl acetate. Stock solutions of the assay compounds were prepared by dissolving each into Milli-Q water and subsequently filter-sterilizing through 0.2-μm membrane Nalgene syringe filters (Thermo Scientific, NY, USA). Liquid cultures were grown from single colonies in marine broth until an OD_600_ ~0.3 was reached. One milliliter aliquots of each culture were then centrifuged at 15,000 rpm (Eppendorf Centrifuge 5424) for 1 min and the pellets were resuspended in 1 mL 10% marine broth diluted with sterile seawater. 5 μL of this bacterial stock were subsequently used to inoculate triplicate tubes containing 100 μM of each molecule in sterile 10% marine broth at a ratio of 1:1000. Growth in 10% marine broth without adding metabolites served as negative control. Absorbance at 600 nm of all cultures was measured every 24 hours from 100 μL aliquots dispensed into 96-well flat-bottom plates using an Epoch microplate spectrophotometer (BioTek Instruments Inc. Winooski, VT, USA). Sterile 10% marine broth was used as blank to correct for background media absorbance. Absorbance readings were normalized against the highest value for each bacterial isolate over the assay period. Significant differences in growth were determined by Student’s *t*-test (*p* < 0.05).

### Bacterial motility assay

Semisolid (0.25% w/v) marine broth agar plates supplemented with a final concentration of 2 μM rosmarinic acid were used to assess its effect on the motility of bacterial strains *S. pseudonitzschiae* F5, *Phaeobacter* sp. F10, and *A. macleodii* F12. Each strain was incubated in marine broth overnight then gently inoculated, using a sterilized toothpick, at the center of the agar surface. Triplicate plates were incubated at 26°C for 3 days, after which the proportion of motility area was measured using the ImageJ software (http://rsb.info.nih.gov/ij/) by calculating the area of bacterial diffusion. Significant differences between control plates and rosmarinic acid-treated plates were determined by Student’s *t*-test (*p*< 0.05).

### Fluorescence microscopy

Axenic *A. glacialis* A3 cultures with an initial cell density of ~4,000 cells/mL in the mid-exponential phase were inoculated with cultures of strains *S. pseudonitzschiae* F5, *Phaeobacter* sp. F10, and *A. macleodii* F12 at a cell density of ~1×10^4^ cells/mL grown overnight in marine broth at 26°C after centrifugation at 4000 rpm for 10 mins followed by washing twice with sterile *f/2* medium. One mL co-cultures of *A. glacialis* A3 with strains *S. pseudonitzschiae* F5, *Phaeobacter* sp. F10, and *A. macleodii* F12 in mid-exponential phase were gently filtered onto 3-μm 25 mm polycarbonate membrane filters (Whatman). 8 μL Moviol-SYBR Green I (Thermo Fisher Scientific, MA) mixture was used to stain cells as described previously (72) and 1 mL Alcian blue in 0.06% glacial acetic acid (pH 2.5) was used to stain transparent exopolymeric particles for 10 min at room temperature. Samples were visualized on an epifluorescent microscope (Leica DMI6000 B, Germany) using L5 and Y5 fluorescence filter sets.

### AzeR global distribution and homology analysis

The amino acid sequence of an azelaic acid transcriptional regulator, AzeR, from Bez *et al* (24) was used to search for potential homologs in the bacterial isolates *S. pseudonitzschiae* F5, *Phaeobacter* sp. F10, and *A. macleodii* F12 on BLASTX (e-value threshold of 1e-05). The resulting hits from the consortium isolates and the AzeR sequence were then used to generate a hidden Markov model (HMM) profile on HMMER v3.1b2 with hmmbuild. The hmm profile was queried against the *Tara* Oceans Microbiome Reference Gene Catalog version 1 on the Ocean Gene Atlas (http://tara-oceans.mio.osupytheas.fr/ocean-gene-atlas/) webserver (114) with an e-value threshold of 1e-50 and a bitscore threshold of 150. Geographical distributions and taxonomic abundances of homologs found in surface and deep chlorophyll maximum samples across all size fractions (0-3 μm) were visualized as donut plots across a world map. The same hmm profile was then queried against the Pfam database (115) with an e-value threshold of 1e-100, resulting in 1,621 hits. Duplicate hits, hits with <200 amino acids, and hits with no taxonomic classification were discarded. The remaining sequences were clustered on USEARCH (116) with an identity threshold of 90%. The resulting 1,043 sequences, in addition to the ones used to build the hmm profile, were aligned on MUSCLE v3.8.31 and the alignment trimmed using trimAl v1.267 (117) on “gappyout” mode. FastTree v2.1.10 (118) was used to infer phylogeny and the unrooted tree was visualized on the Interactive Tree of Life (iTOL) tool v5 (119).

### Data deposition and materials availability

The *Asterionellopsis glacialis* strain A3 is available from the National Center for Marine Algae and Microbiota (NCMA) collection under the accession CCMP3542. The *A. glacialis* A3 genome is deposited at DDBJ/ENA/GenBank under the accession WKLE01000000 in NCBI-BioProject PRJNA588343. RNA-seq reads of *A. glacialis* A3 are deposited in NCBI under the BioProject number PRJNA588343. Metagenomic reads and RNA-seq reads of the bacterial consortium are deposited in NCBI under the BioProject number PRJNA578578. Metagenomically-assembled genomes are deposited at DDBJ/ENA/GenBank under the accessions WKFI01000000-WKFN01000000 in NCBI-BioProject PRJNA588964. Whole genome assemblies of consortium-isolated strains *Sulfitobacter pseudonitzschiae* F5, *Phaeobacter* sp. F10, and *Alteromonas macleodii* F12 are deposited at DDBJ/ENA/GenBank under the accessions WKFG01000000, WKFH01000000, and CP046140-CP046144, respectively in NCBI-BioProject PRJNA588972. All mass spectral datasets are deposited in the MassIVE database (https://massive.ucsd.edu) under accession MSV000084592. All software packages used in this study are free and open source.

## Supporting information

AzeR HMMprofile

## Acknowledgments

We thank the NYU Abu Dhabi Core Technology Platforms for support related to genomics sequencing and mass spectrometry. We also thank Dain McParland for help collecting water samples, Bryndan P. Durham and Elodie Ghedin for helpful comments on the manuscript. This project was supported by NYU Abu Dhabi Research Fund AD179 to S.A.A.

## Author Contributions

S.A.A., G.B., M.A.O. and C.R.V. designed experiments. G.B. carried out the reseeding experiment and extracted DNA and RNA. M.A. prepared sequencing libraries. M.A.O. ran and processed metabolomic samples. C.F. and A.I. carried out the bacterial isolation, microscopy and growth experiments. A.A.S., A.C., N.D., K.C.G. and M.P.S. developed and carried out supporting algorithms, bioinformatic analyses and computational pipelines. A.A.S., M.A.O., A.C., C.F., A.I., and S.A.A. analyzed the data. A.A.S. and S.A.A. wrote the manuscript with input from all coauthors.

The authors declare no competing interests.

---

**Figure S1.**
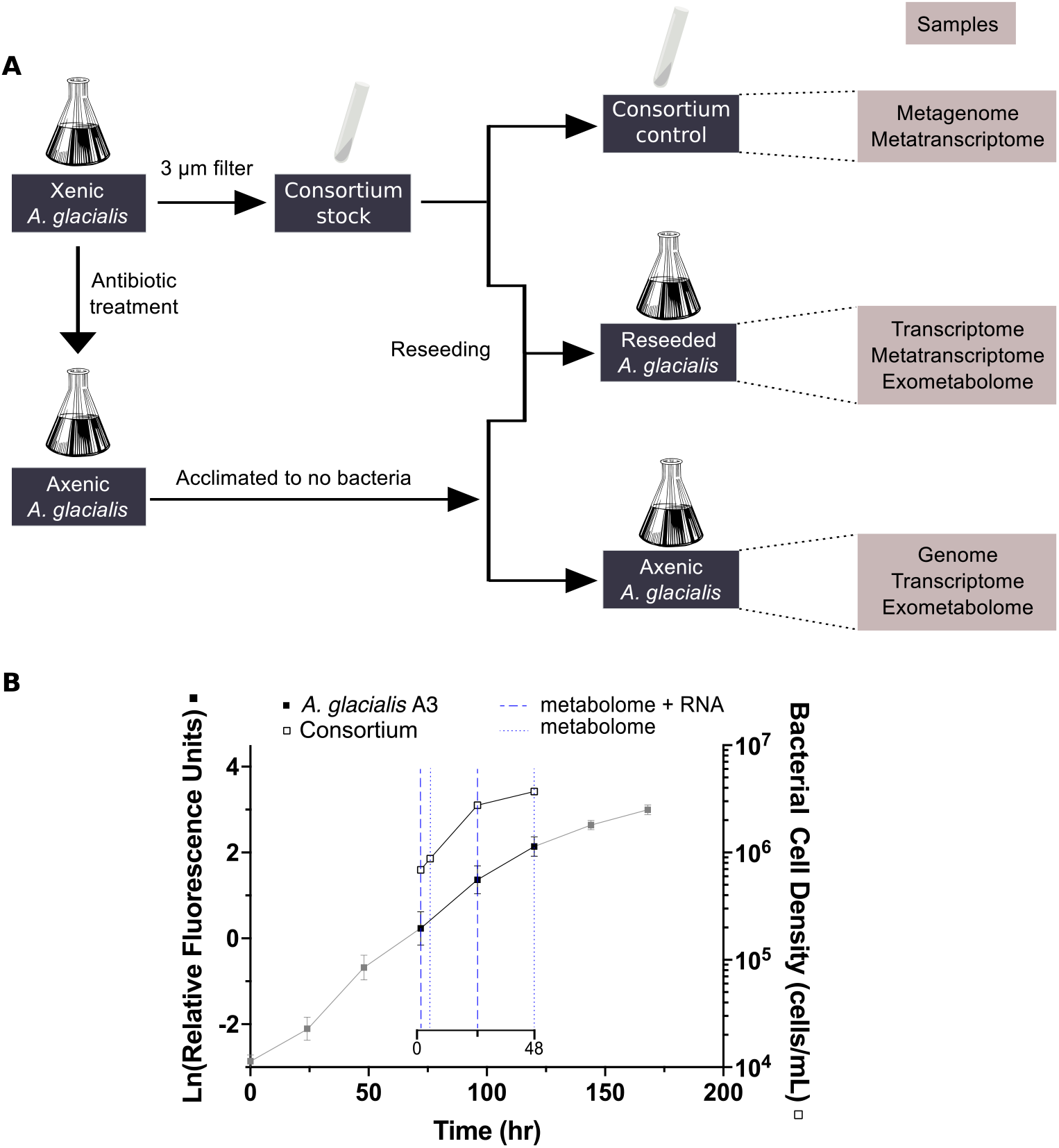
Experimental schematic and growth of the diatom and bacterial consortium during reseeding. (**A**) Scheme of the reseeding experiment described in the Methods. Briefly, xenic *A. glacialis* A3 culture was made axenic using antibiotics and the resulting axenic culture was acclimated to absence of bacteria and subsequently used for genome sequencing. To reseed this axenic culture with bacteria, xenic *A. glacialis* A3 cultures were used to remove diatom cells and obtain a consortium stock, a portion of which was used to obtain a consortium metagenome. At the beginning of the reseeding experiment, the consortium stock was added to either axenic *A. glacialis* A3 or to sterile media (for the consortium RNA negative control) as described in the Methods. A second axenic *A. glacialis* A3 culture served as diatom control. RNA and exometabolomes were collected at different time points from each set of samples. (**B**) Growth of *A. glacialis* A3 and the microbial consortium. Because all cultures were grown in seawater-based *f/2* media that does not support significant heterotrophic growth, bacterial growth after reintroducing the microbial consortium to the diatom indicated uptake of diatom-excreted DOM. Closed squares represent diatom *in vivo* chlorophyll *a* fluorescence while open squares represent bacterial cell density. Grey points on the *A. glacialis* A3 growth curve denote time points before and after sampling. The secondary x-axis indicates the beginning of reseeding of the consortium (t=0). Dashed lines indicate time points at which RNA and metabolome samples were collected (0.5, 24 hours) while dotted lines indicate time points at which only metabolome samples were collected (4, 48 hours). Error bars represent standard deviation (SD) of triplicate cultures.

**Figure S2.**
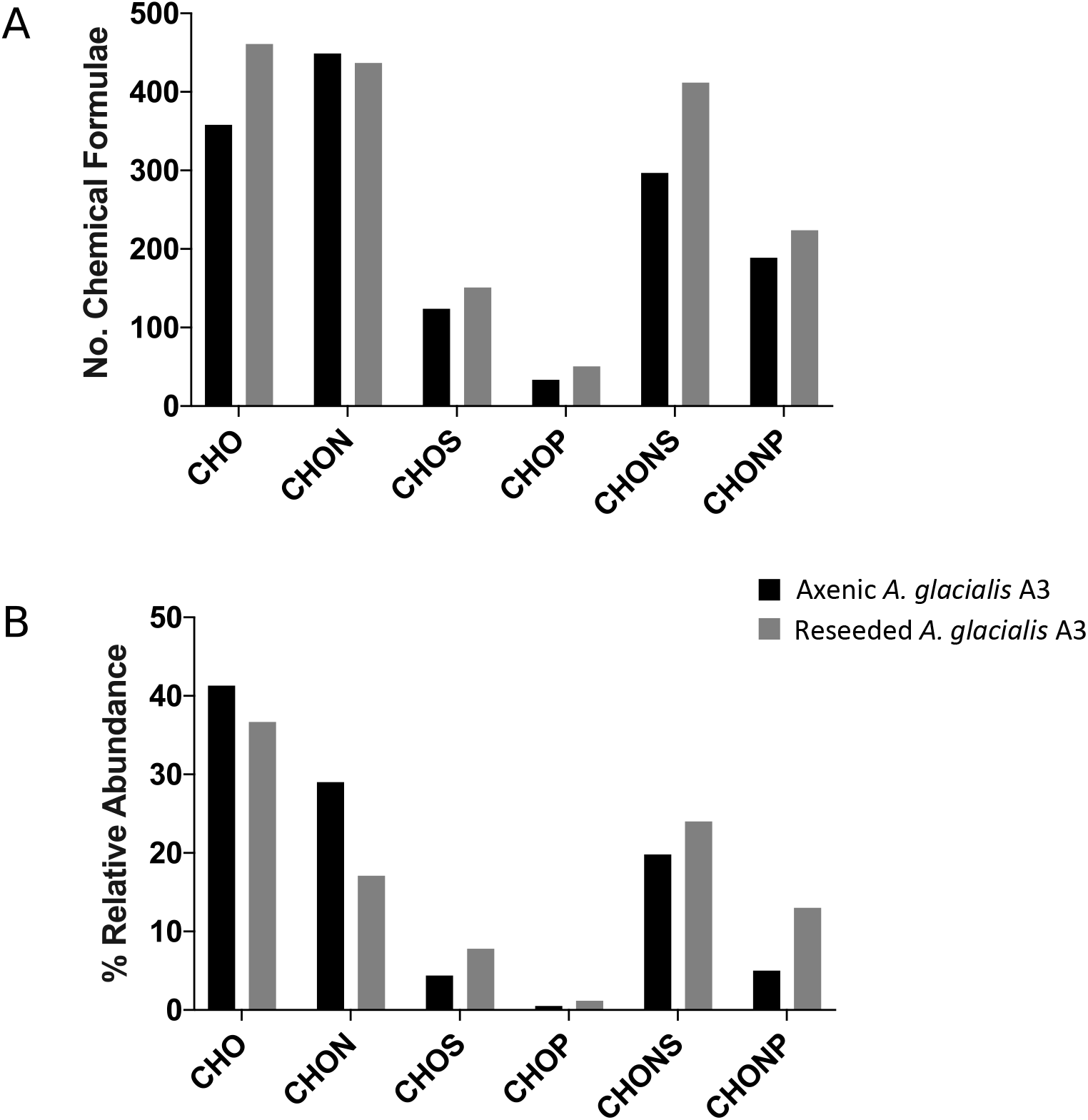
Diversity and abundance of chemical formulae of SPE-extracted DOM in the exometabolome. Exometabolomes were analyzed on a FT-ICR-MS as described in the Methods. **(A)** Number of SPE-extracted chemical formulae from the axenic diatom and reseeded cultures 24 hours after reseeding. (**B)** Relative abundance of chemical formulae in the axenic diatom and reseeded cultures. Chemical formulae calculation was performed with an error threshold of 0.5 ppm from the exact mass for each chemical formula and isotopic fine structure. Chemical formulae were only included in the analysis if 100% of theoretical isotope peaks matched the isotopic fine structure of each formula. Biological samples were pooled to acquire sufficient signal for analysis.

**Figure S3.**
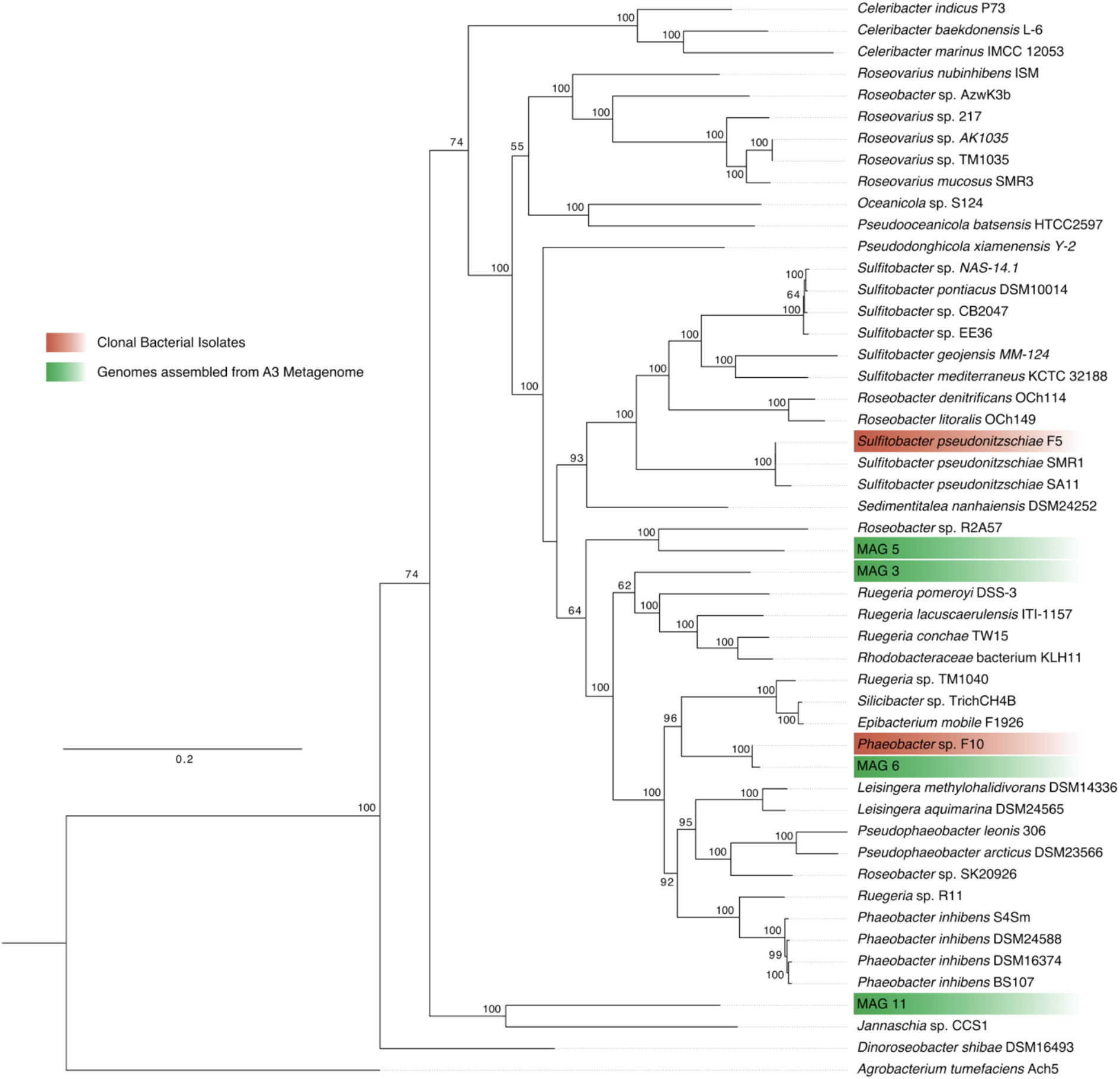
Metagenomically-assembled genomes (MAGs) and isolated bacterial strains from the consortium belonging to roseobacters. Maximum-likelihood phylogeny of whole-genome sequences from 50 bacterial strains comprising 43 species from the Rhodobacteraceae family, four MAGs, two isolates, and one outgroup species *(Agrobacterium tumefaciens* Ach5). Numbers adjacent to branches represent node support calculated with 1,000 bootstraps. Accession numbers of all strains are listed in Table S6.

**Figure S4.**
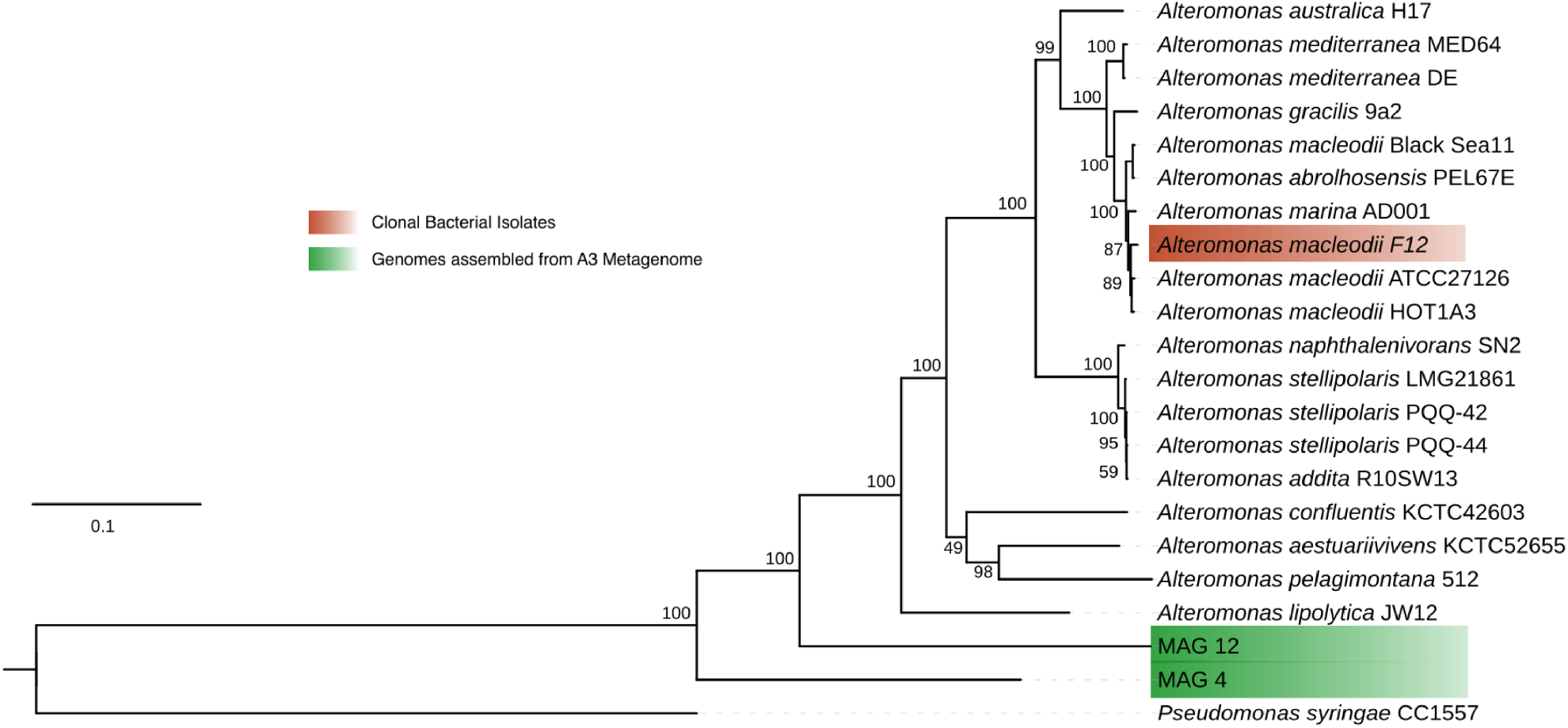
MAGs and isolated bacterial strains from the consortium belonging to Alteromonadaceae. Maximum-likelihood phylogeny of whole-genome sequences from 22 bacterial strains comprising 18 species from the Alteromonadaceae family, two MAGs, one isolate, and one outgroup species *(Pseudomonas syringae* CC1557). Numbers adjacent to branches represent node supports calculated with 1,000 bootstraps. Accession numbers of all strains are listed in Table S7.

**Figure S5.**
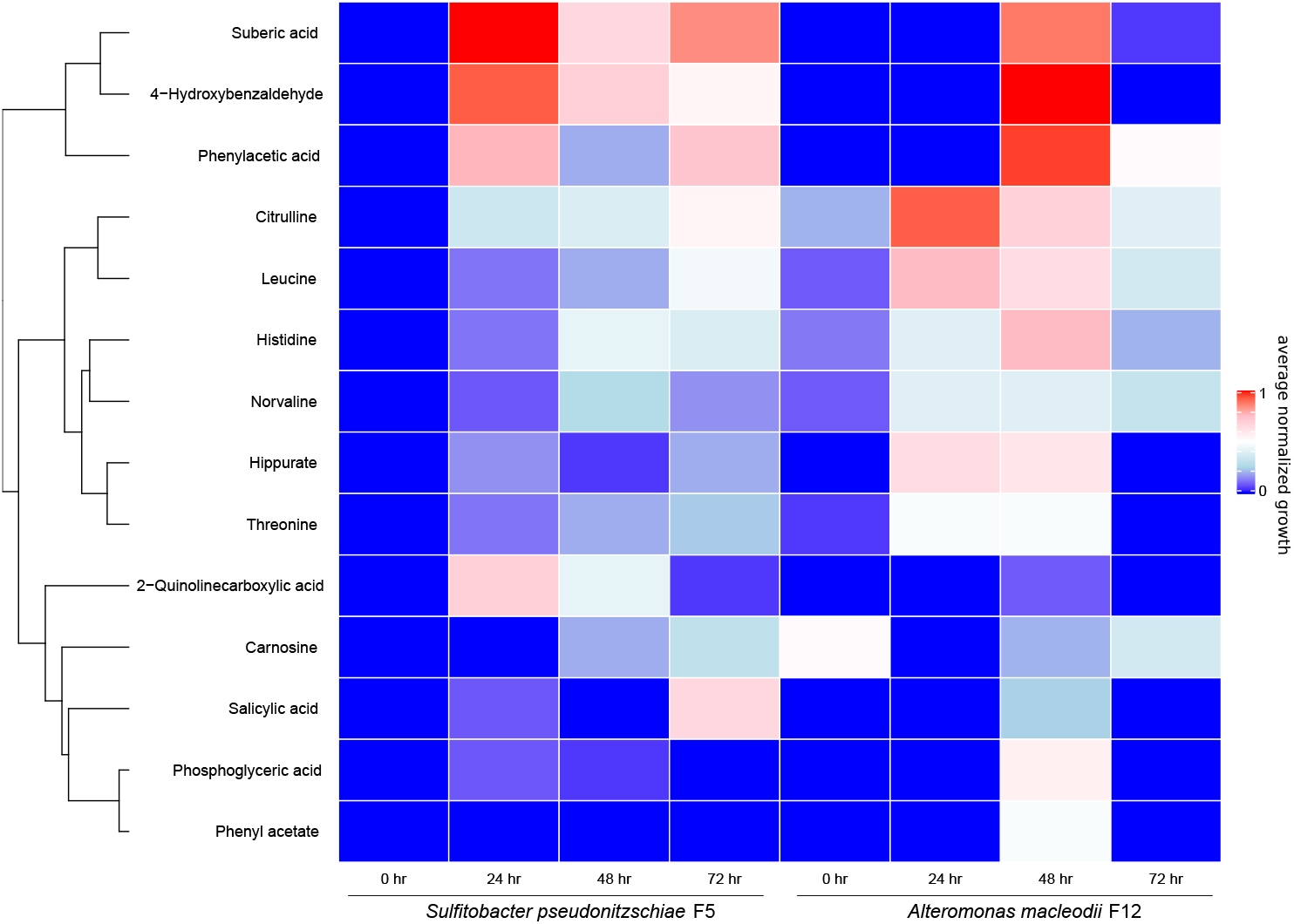
A subset of confirmed metabolites promotes the growth of strains *S. pseudonitzschiae* F5 and *A. macleodii* F12 isolated from the *A. glacialis* A3 consortium. Cell density (OD_600_) of *S. pseudonitzschiae* F5 and *A. macleodii* F12 grown on a subset of confirmed metabolites from the exometabolome. Bacteria were grown in 10% marine broth supplemented with 100 μM final concentration of each molecule. Colors represent average growth from biological triplicates normalized to growth on 10% MB control.

**Figure S6.**
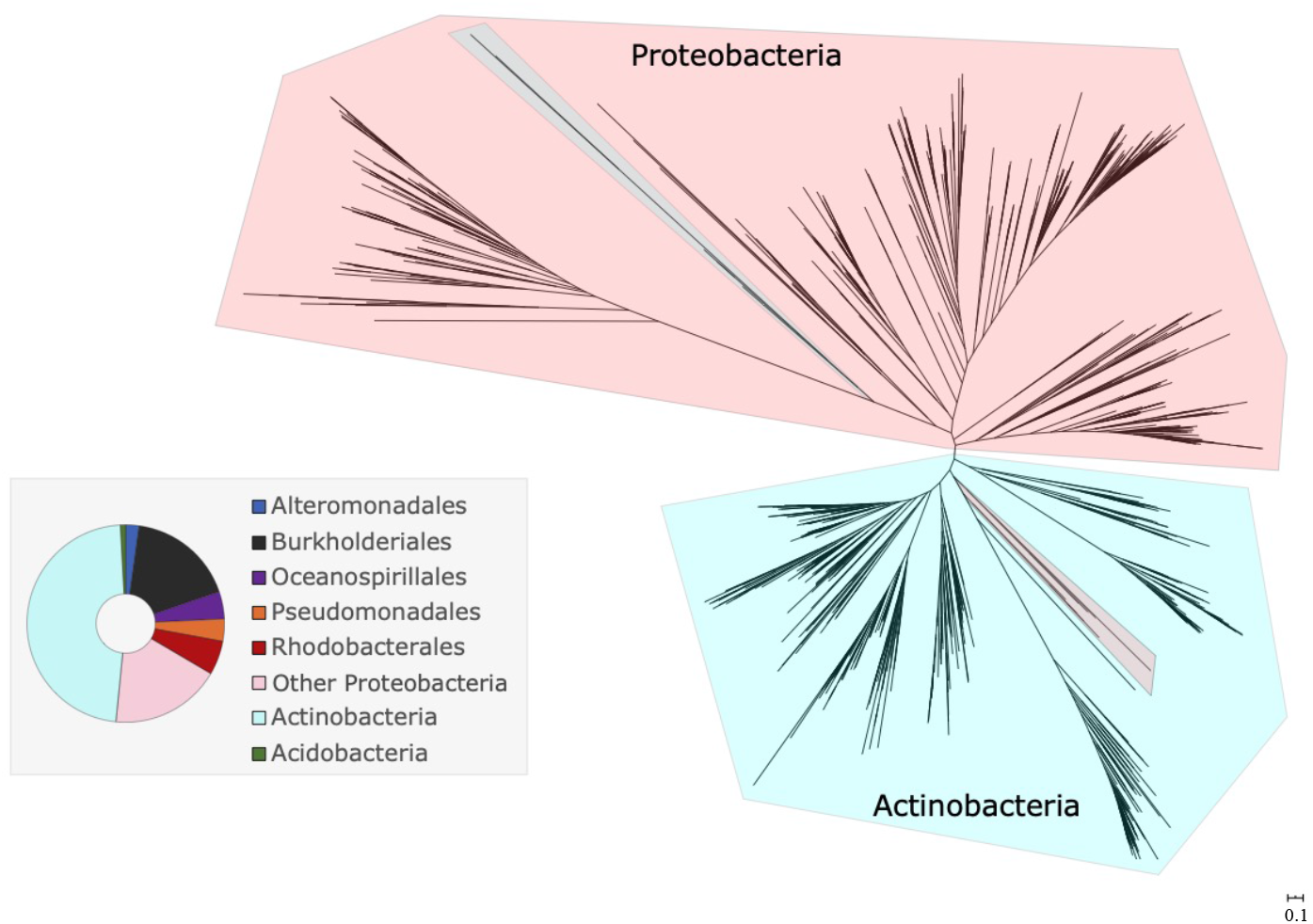
Bacterial response to azelaic acid is restricted to a handful of taxa. Maximum-likelihood tree of the azelaic acid transcriptional regulator, AzeR, from 1,043 protein sequences shows that response to azelaic acid in the Proteobacteria phylum is restricted to mostly five taxa. The donut plot depicts the taxonomy of all homologs.

**Table S1.** Summary of the assembly of metagenomically-assembled genomes (MAGs) recovered from the microbial consortium shotgun metagenome and their closest reference genome from the NCBI Reference Sequence Database according to amino acid identity (AAI).

| MAG | Recruitment in bacterial consortium metagenome (%) | Total length (bp) | Contigs | Predicted genes | N50 | GC content (%) | Completeness (%) | Closest genome in RefSeq | AAI (%) |
| --- | --- | --- | --- | --- | --- | --- | --- | --- | --- |
| 3 | 0.26 | 4761801 | 26 | 4515 | 499224 | 58.70 | 99.49 | *Phaeobacter gallaeciensis* | 67.38 |
| 4 | 3.99 | 4662739 | 66 | 4310 | 109594 | 43.57 | 91.67 | *Alteromonas australica* | 57.7 |
| 5 | 0.44 | 3964870 | 16 | 3851 | 483383 | 57.71 | 99.47 | *Phaeobacter gallaeciensis* | 63.58 |
| 6 | 9.35 | 3933228 | 60 | 3850 | 150332 | 60.01 | 98.58 | *Phaeobacter gallaeciensis* | 86.2 |
| 8 | 2.07 | 3505326 | 20 | 3269 | 267358 | 44.08 | 99.14 | *Thalassolituus oleivorans*<br>MIL 1 | 51.93 |
| 9 | 34.18 | 3330214 | 59 | 3031 | 78673 | 34.53 | 98.49 | *Polaribacter reichenbachii* | 64.59 |
| 10 | 8.31 | 2639318 | 3 | 2602 | 1552570 | 57.02 | 99.2 | *Altererythrobacter ishigakiensis* | 96.99 |
| 11 | 1.46 | 2548972 | 12 | 2529 | 306111 | 56.15 | 98.79 | *Roseibacterium elongatum*<br>DSM 19469 | 62.11 |
| 12 | 1.03 | 2510135 | 16 | 2343 | 188996 | 43.72 | 96.03 | *Alteromonas australica* | 60.34 |
| 13 | 14.64 | 2198323 | 12 | 2271 | 368442 | 48.93 | 94.88 | *Halomonas aestuarii* | 48.42 |

**Table S2.** Average % mRNA reads mapped to the MAGs relative to total mRNA reads after quality control and % DE genes for MAGs relative to total number of genes in each MAG.

|  | Consortium only at 0.5 hours | Reseeded consortium at 0.5 hours |  | Reseeded consortium at 24 hours |  |
| --- | --- | --- | --- | --- | --- |
| MAG | average % mRNA mapped | average % mRNA mapped | DE genes (%) | average % mRNA mapped | DE genes (%) |
| **3** | 0.85 | 1.05 | 14.05 | 0.51 | 1.08 |
| **4** | 6.26 | 0.58 | 0.16 | 0.69 | 1.61 |
| **5** | 0.37 | 0.56 | 11.76 | 0.58 | 0 |
| **6** | 9.63 | 30.13 | 4.97 | 24.92 | 5.84 |
| **8** | 0.71 | 1.74 | 0.09 | 6.08 | 0 |
| **9** | 42.97 | 3.86 | 2.91 | 3.12 | 7.54 |
| **10** | 7.81 | 16.99 | 4.19 | 4.02 | 6.07 |
| **11** | 0.53 | 2.41 | 6.73 | 0.69 | 0 |
| **12** | 1.56 | 0.56 | 2.39 | 1.15 | 0.04 |
| **13** | 13.84 | 2.73 | 2.38 | 0.85 | 3.40 |

**Table S3.**
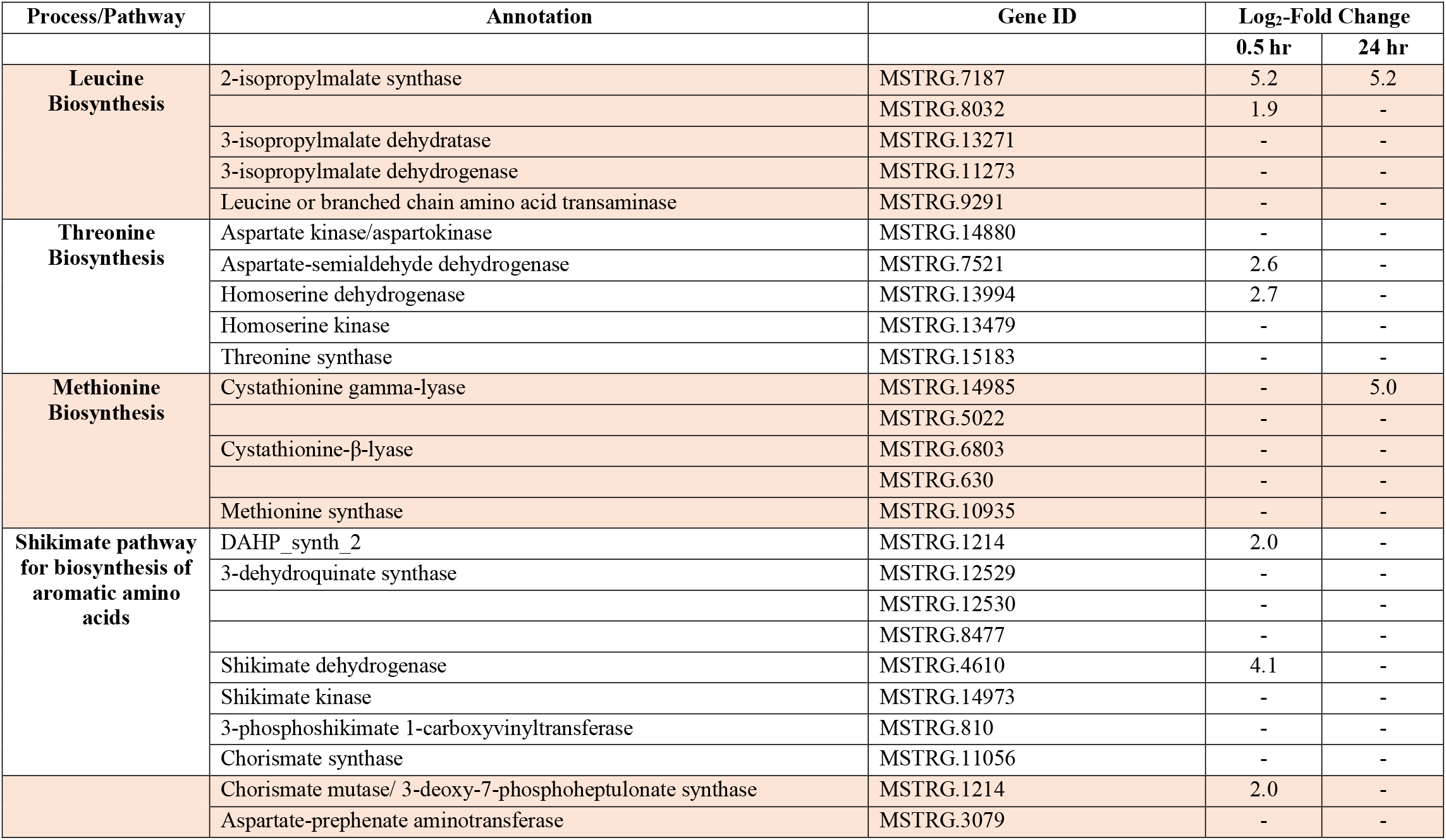
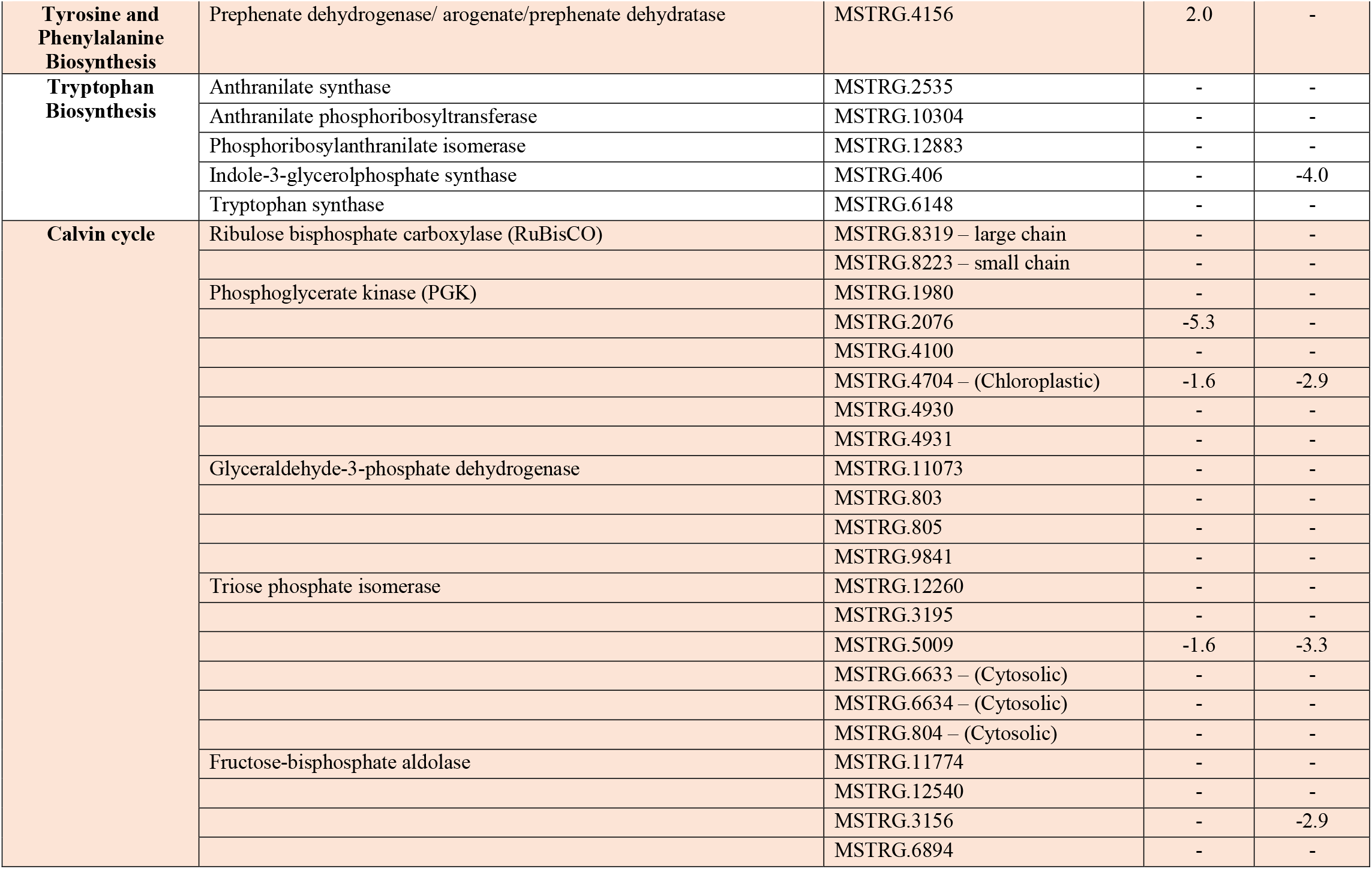
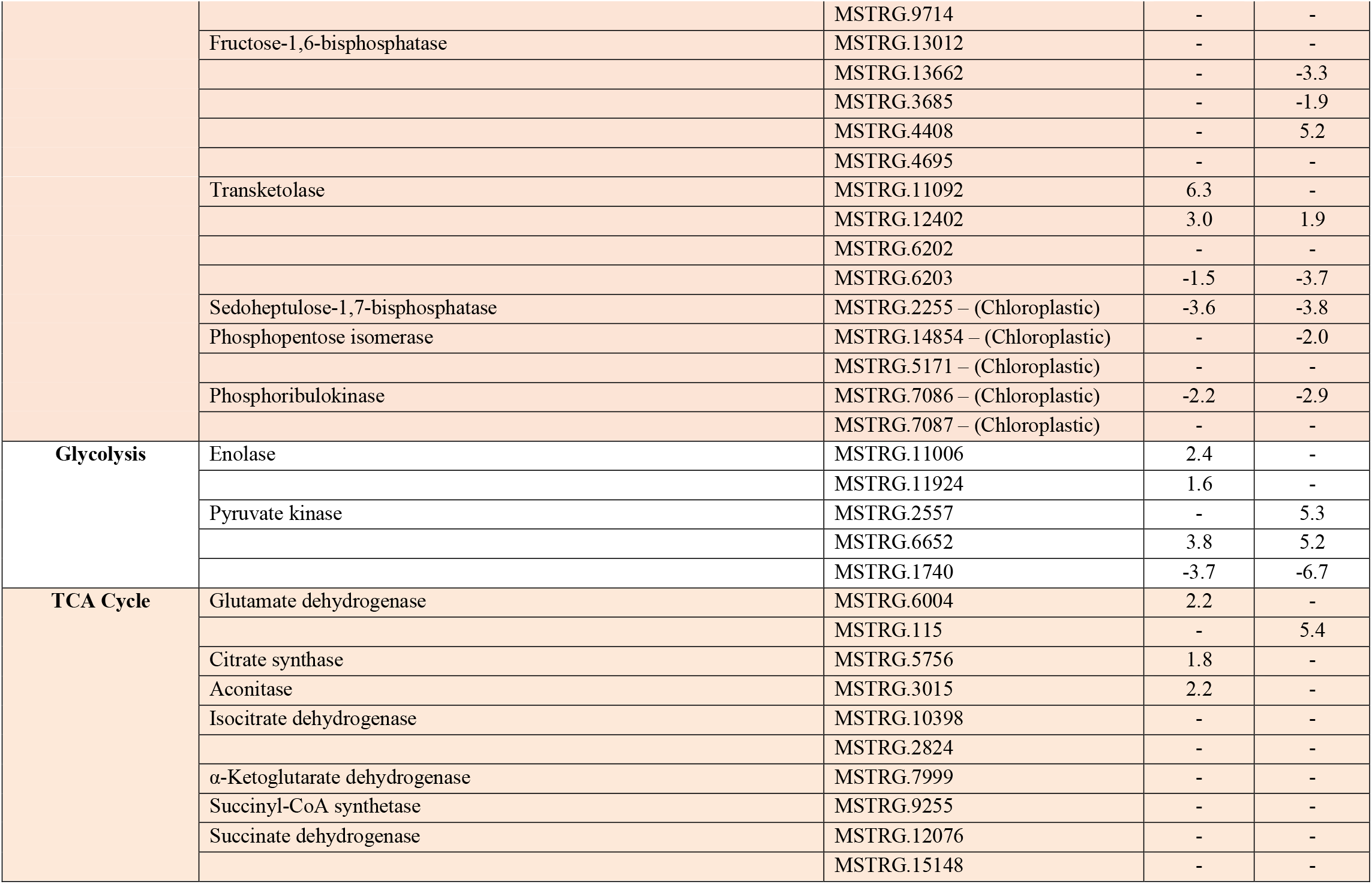
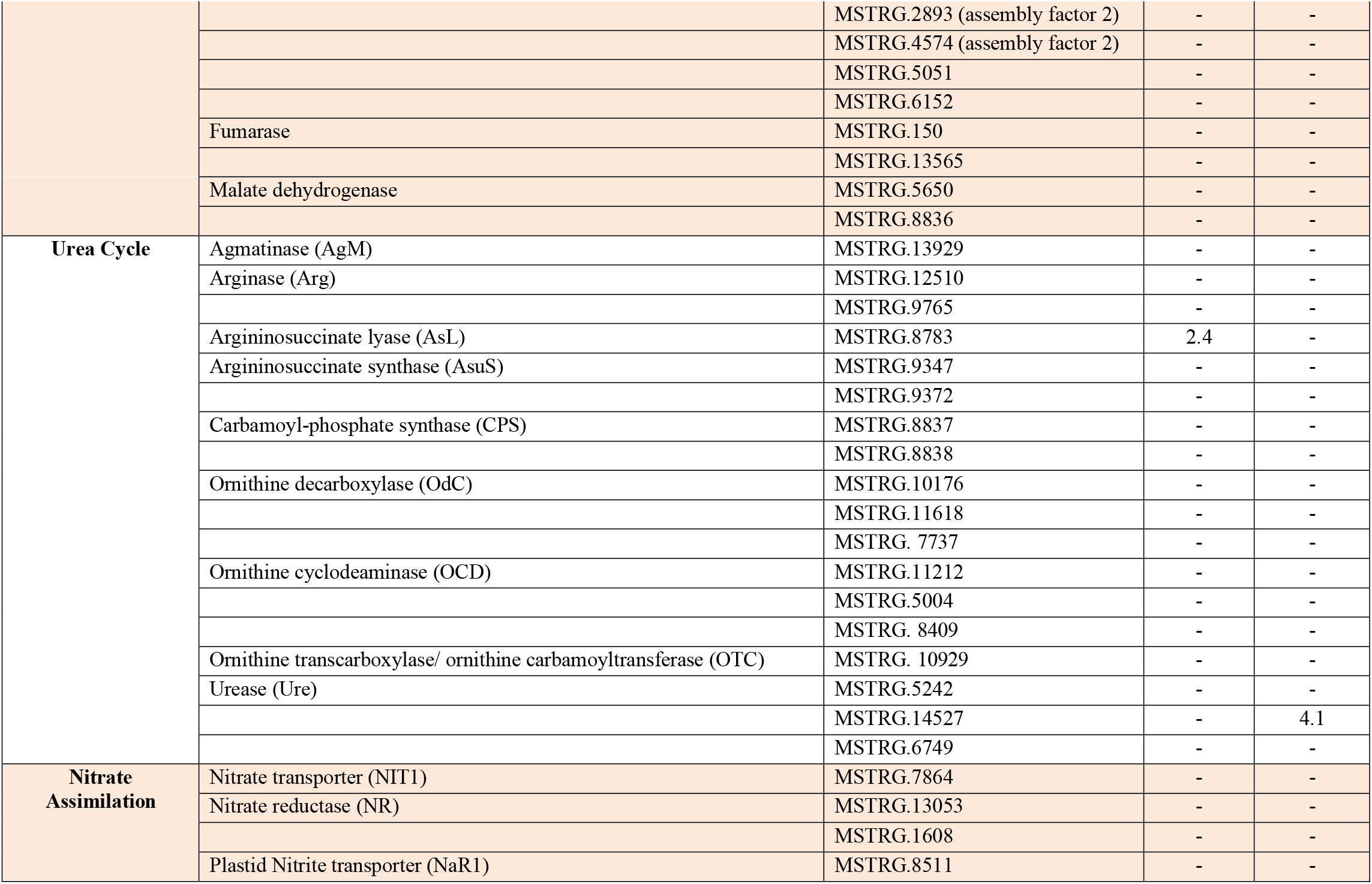
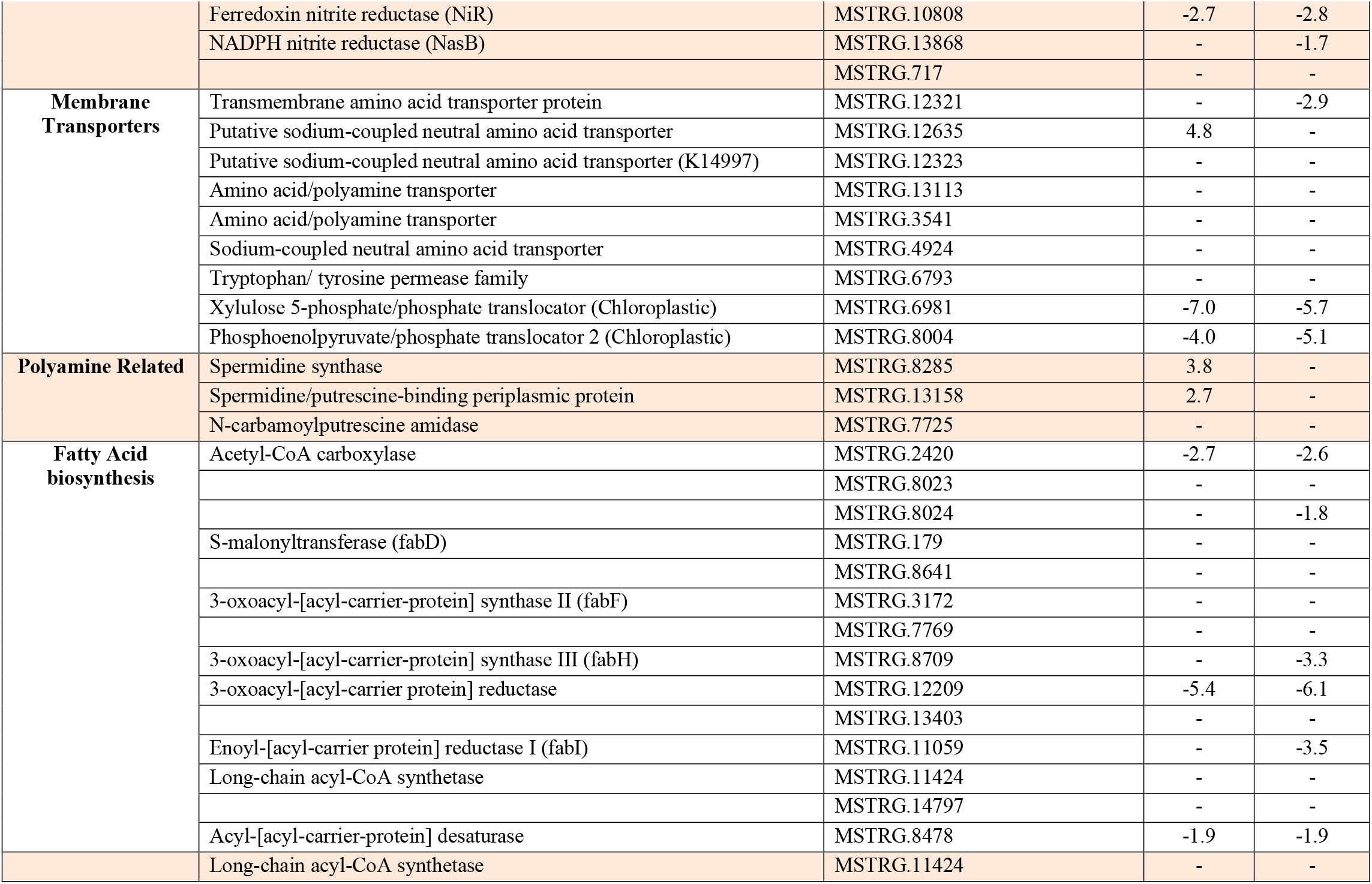
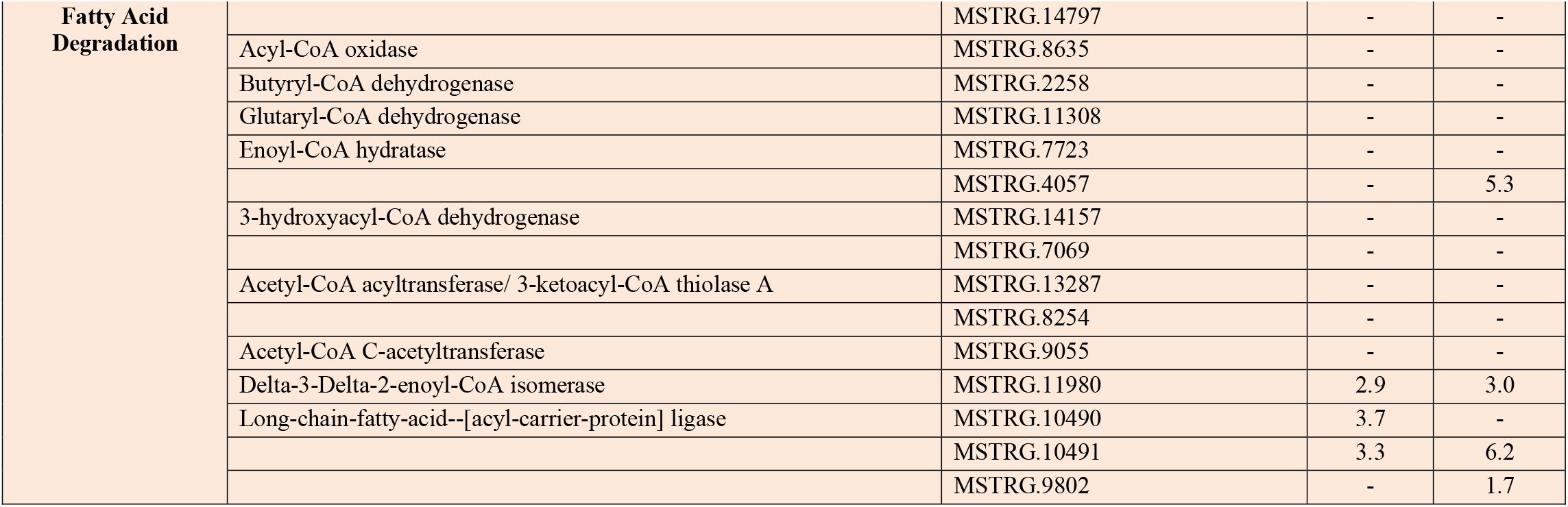
List of selected expressed genes in *A. glacialis* A3, including genes depicted in Fig. 3. Genes with a false discovery rate (FDR) adjusted *p*-value < 0.1 were considered to be differentially expressed. The values correspond to the log_2_-fold change at the two timepoints in response to reseeding relative to axenic controls. Blank cells indicate no differential expression.

**Table S4.** List of confirmed metabolites in Fig. 2*D*. Metabolites were confirmed by comparing retention time, accurate mass, isotopic pattern and fragmentation pattern of each metabolite to a library of in-house chemicals (Mass Spectrometry Metabolite Library of Standards, IROA Technologies, US). Additional molecules were confirmed using the Bruker MetaboBASE Plant Library and MetaboBASE Personal Library 2.0 (BrukerDaltonik, Germany). We were not able to confirm the annotations of molecules found only in reseeded samples. Analysis was done using Metaboscape v4.0 (BrukerDaltonik, Germany). Primary Database refers to the in-house chemical library. M= monoisotopic mass, RT= retention time, ΔmDA= mass difference in milliDalton, IP= isotopic pattern, Δppm= mass difference in parts per million.

| M (Da) | RT (min) | $\Delta mDa$ | IP | MS/MS Score | Chemical Formula | Adduct | Name | $\Delta ppm$ | Database |
| --- | --- | --- | --- | --- | --- | --- | --- | --- | --- |
| 103.0622 | 19.94 | 0.76 | 80 | 980.1 | C4H9NO2 | [4M+H]<sup>+</sup> | 4-Aminobutanoate | 7.4 | Primary Database |
| 117.078 | 0.45 | 1 | 39 | 999.9 | C5H11NO2 | [M+H+H]<sup>2+</sup> | Norvaline | 8.5 | Primary Database |
| 119.0578 | 12.42 | 0.22 | 46 | 760.9 | C3H7NO3 | [M+H]<sup>+</sup> | Threonine | 1.8 | Primary Database |
| 122.0369 | 10.25 | 0.06 | 45 | 773.6 | C7H6O2 | [M+H]<sup>+</sup> | 4-Hydroxybenzaldehyde | 0.5 | Primary Database |
| 131.0946 | 13.51 | 0.39 | 10 | 761.6 | C6H13NO2 | [M+H]<sup>+</sup> | Leucine | 3 | Primary Database |
| 136.0511 | 19.91 | 0.9 | 62 | 967.5 | C8H8O2 | [M+H]<sup>+</sup> | Phenylacetic acid | 6.6 | Primary Database |
| 136.0512 | 11.34 | 0.8 | 52 | 735.3 | C8H8O2 | [M-H]<sup>-</sup> | Phenyl acetate | 5.9 | Primary Database |
| 138.031 | 4.26 | 0.97 | 36 | 922.2 | C7H7NO2 | [M+H]<sup>+</sup> | Salicylic acid | 7 | Primary Database |
| 145.0739 | 8.71 | 0.11 | 23 | 759.1 | C6H11NO3 | [M+H]<sup>+</sup> | 4-Acetamidobutanoic acid | 0.8 | Primary Database |
| 146.0573 | 19.95 | 0.68 | 51 | 993.1 | C6H10O4 | [M+H]<sup>+</sup> | 3-Methylglutaric acid | 4.7 | Primary Database |
| 152.0464 | 18.8 | 0.6 | 59 | 846.5 | C8H8O3 | [M-H]<sup>-</sup> | 4-Hydroxy-phenylacetate | 3.9 | Primary Database |
| 161.0695 | 18.7 | -0.52 | 52 | 860.1 | C6H11NO4 | [M+H]<sup>+</sup> | α-Aminoadipate | -3.2 | Primary Database |
| 167.0577 | 0.38 | 0.33 | 41 | 716.4 | C8H9NO3 | [M-H]<sup>-</sup> | 4-Hydroxy-phenylglycine | 2 | Primary Database |
| 169.0843 | 4.47 | 0.73 | 46 | 949.1 | C7H11N3O2 | [M+H]<sup>+</sup> | 1-Methylhistidine | 4.3 | Primary Database |
| 173.0482 | 1.34 | -0.16 | 41 | 845 | C10H7NO2 | [M+H]<sup>+</sup> | 2-Quinolinecarboxylic acid | -0.9 | Primary Database |
| 174.088 | 0.38 | 1 | 50 | 612 | C8H14O4 | [M+H]<sup>+</sup> | Suberic acid | 5.7 | Primary Database |
| 175.0958 | 6.8 | 0.21 | 70 | 897.9 | C6H13N3O3 | [M+H]<sup>+</sup> | Citrulline | 1.2 | Primary Database |
| 179.0571 | 16.74 | 0.95 | 52 | 989.3 | C9H9NO3 | [M+H]<sup>+</sup> | Hippurate | 5.3 | Primary Database |
| 185.9925 | 9.33 | 0.49 | 56 | 882.7 | C3H7O7P | [M+H]<sup>+</sup> | 3-Phosphoglyceric acid | 2.6 | Primary Database |
| 188.1047 | 6.87 | 0.26 | 70 | 872.7 | C9H16O4 | [M+H]<sup>+</sup> | Azelaic acid | 1.4 | Primary Database |
| 221.0898 | 8.04 | 0.18 | 5.7 | 763.4 | C8H15NO6 | [M+H]<sup>+</sup> | N-acetyl-galactosamine | 0.8 | Primary Database |
| 226.1063 | 11.44 | 0.67 | 33 | 998 | C9H14N4O3 | [M-H]<sup>-</sup> | Carnosine | 3 | Primary Database |
| 282.2556 | 17.01 | 0.4 | 38.7 | 978 | C18H34O2 | [M-H]<sup>-</sup> | Oleic acid | 1.4 | Bruker MetaboBASE Plant Library |
| 284.2709 | 18.17 | 0.61 | 2.9 | 997.6 | C18H36O2 | [M-H]- | Stearic acid | 2.2 | Bruker MetaboBASE<br>Plant Library |
| 301.298 | 10.46 | 0 | 19 | 824.1 | C18H39NO2 | [M+H]+ | Sphinganine | 0 | Primary Database |
| 342.2278 | 15.5 | 0.26 | 40 | 730.5 | C11H14O2 | [M+H]+ | N-Tetradecanoylaspartic acid | 0.8 | Bruker MetaboBASE<br>Personal Library 2.0 |
| 360.0843 | 18.77 | 0.74 | 26 | 928.6 | C18H16O8 | [M+H]+ | Rosmarinic acid | 2.1 | Primary Database |
| 541.3372 | 16.6 | 0.79 | 27 | 992.3 | C25H52NO9P | [M-H]- | C<sub>16</sub>-hydroxy-glycerophosphocholine | 1.5 | Bruker MetaboBASE<br>Plant Library |

**Table S5.**
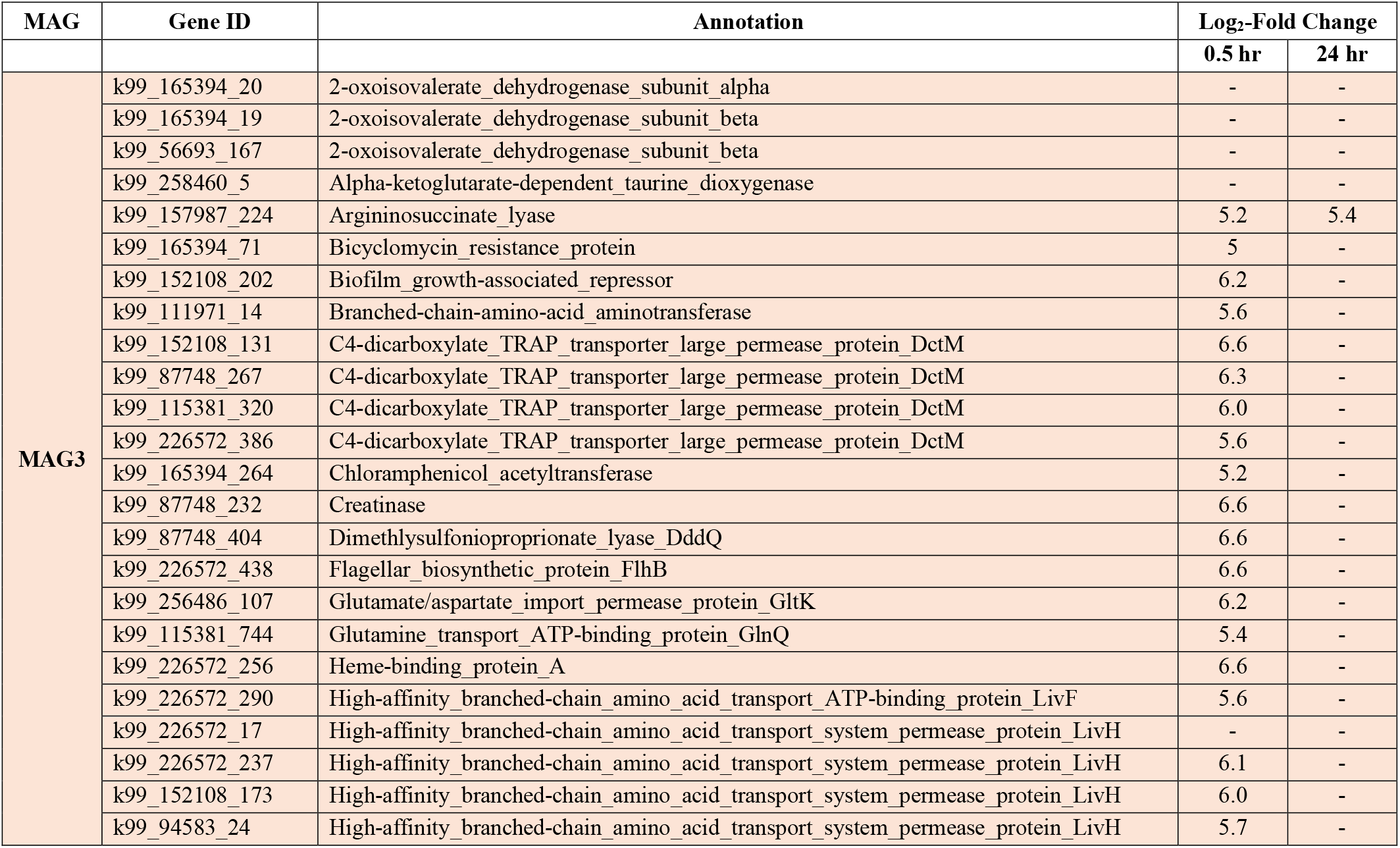
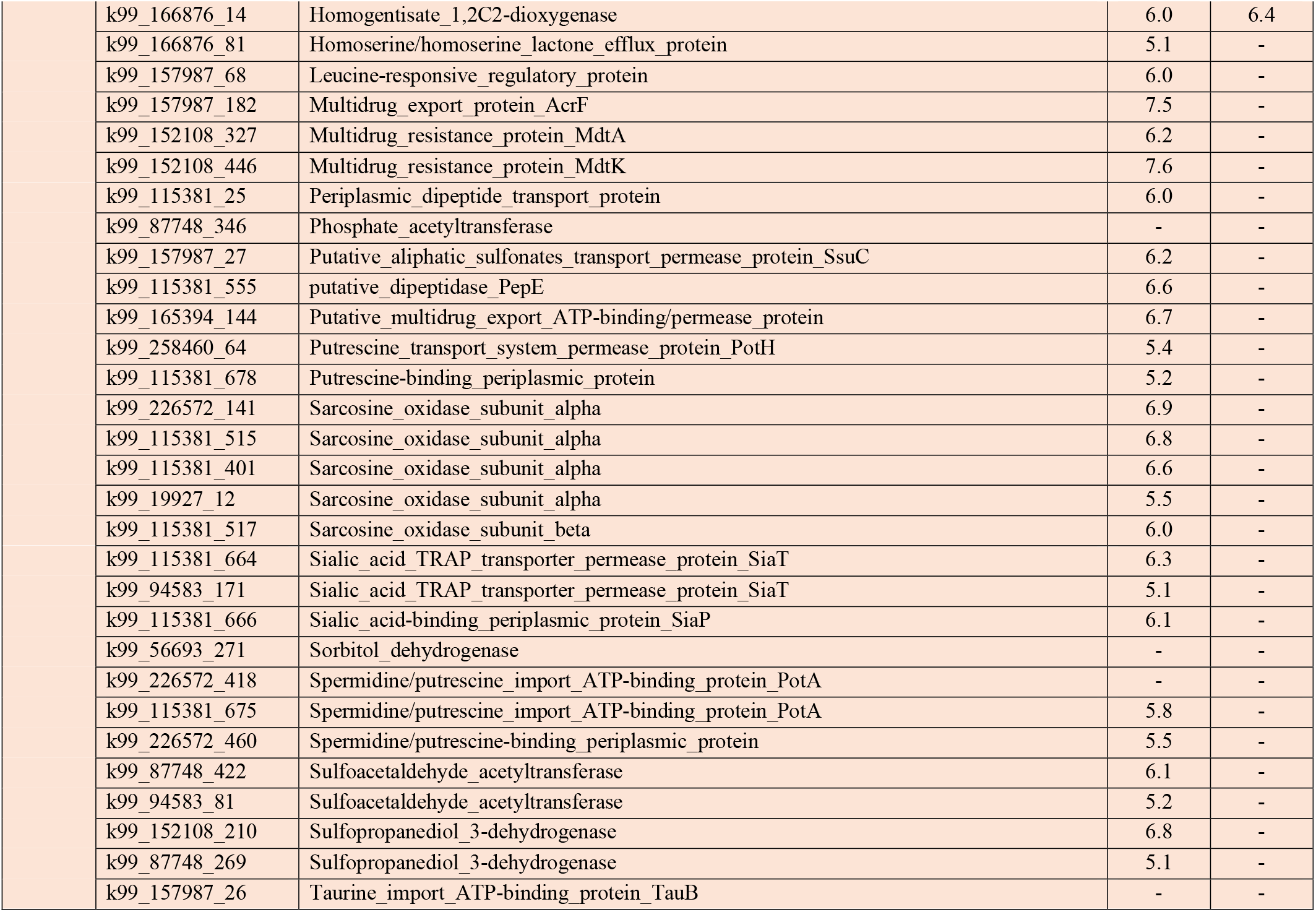
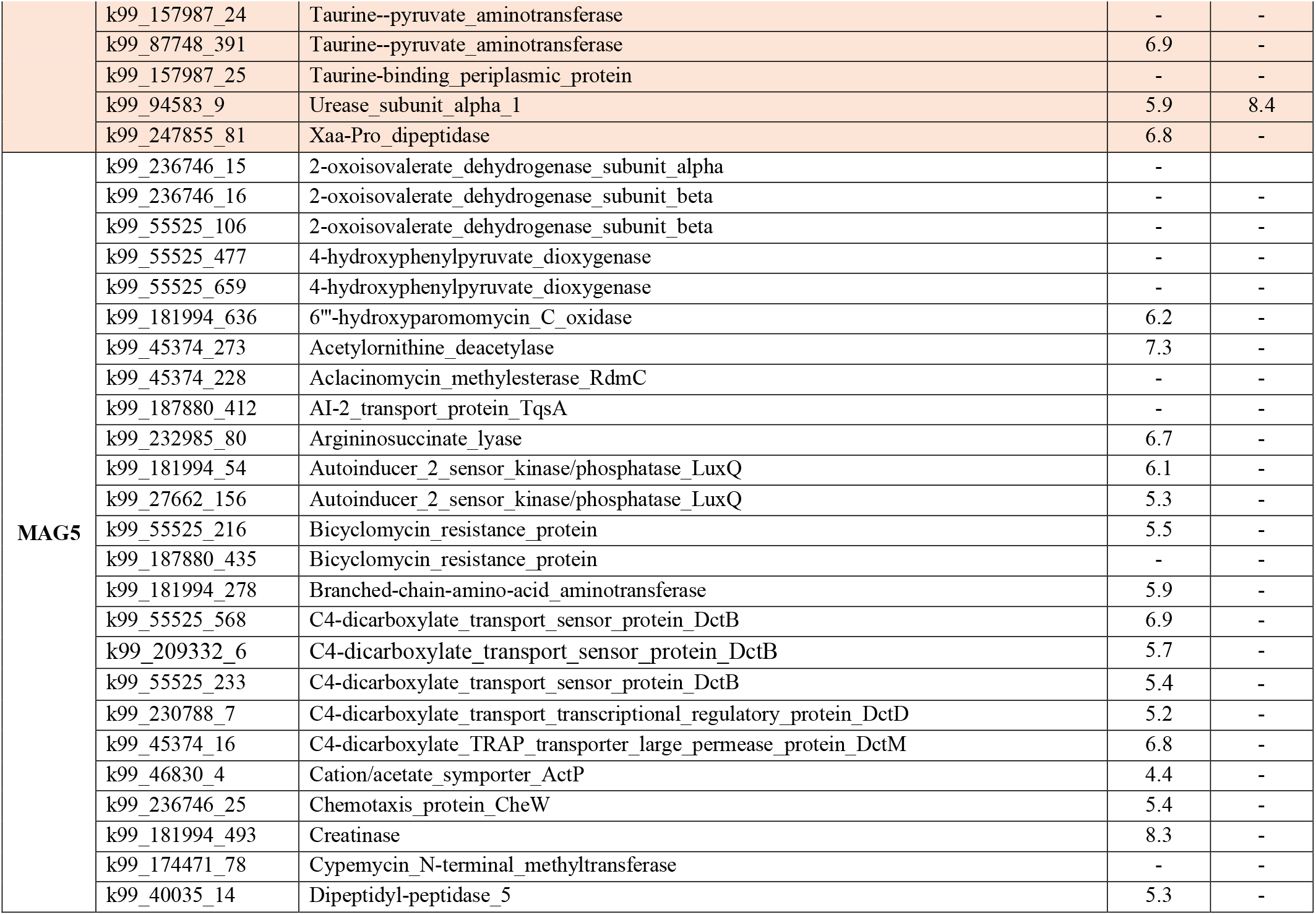
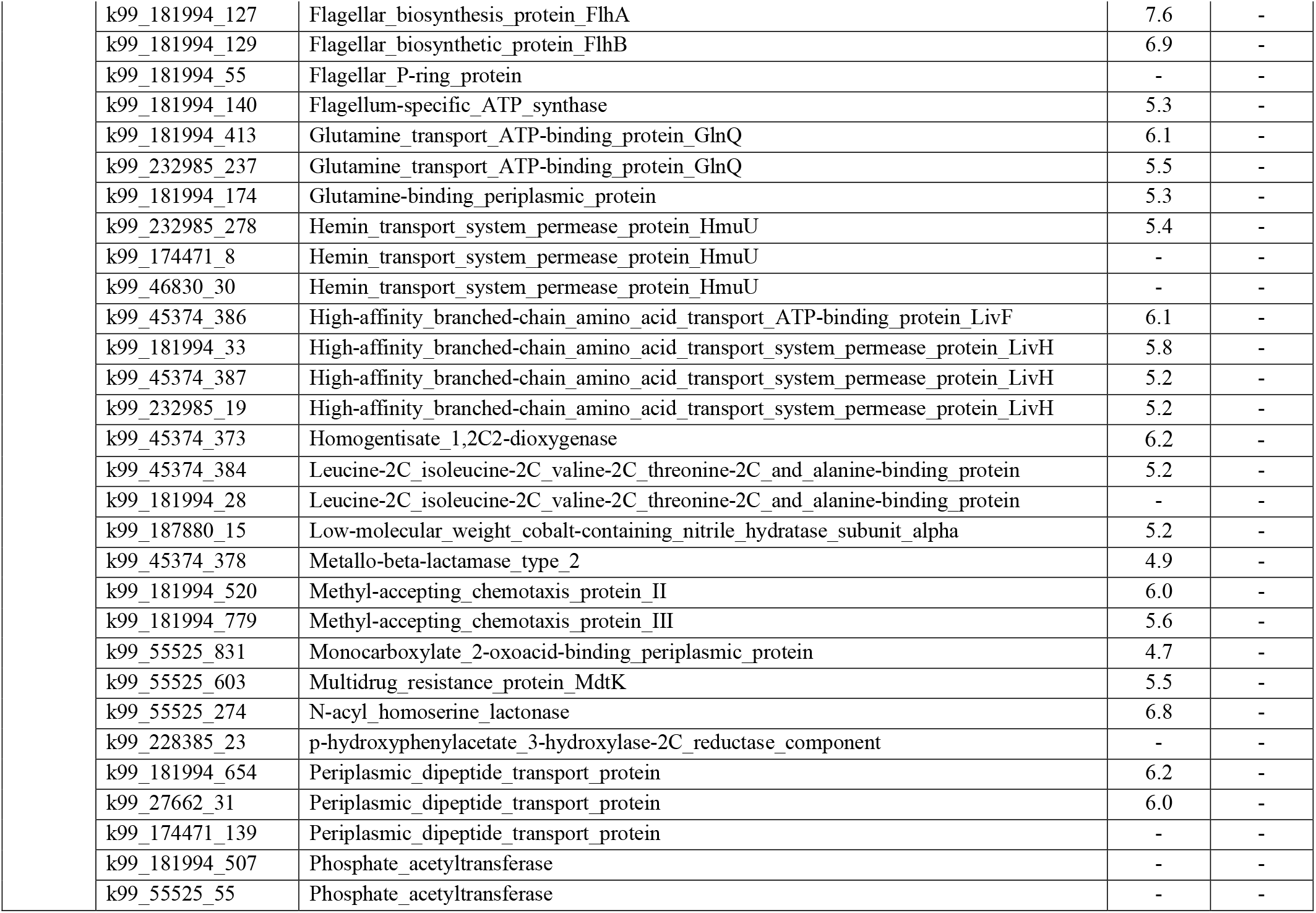
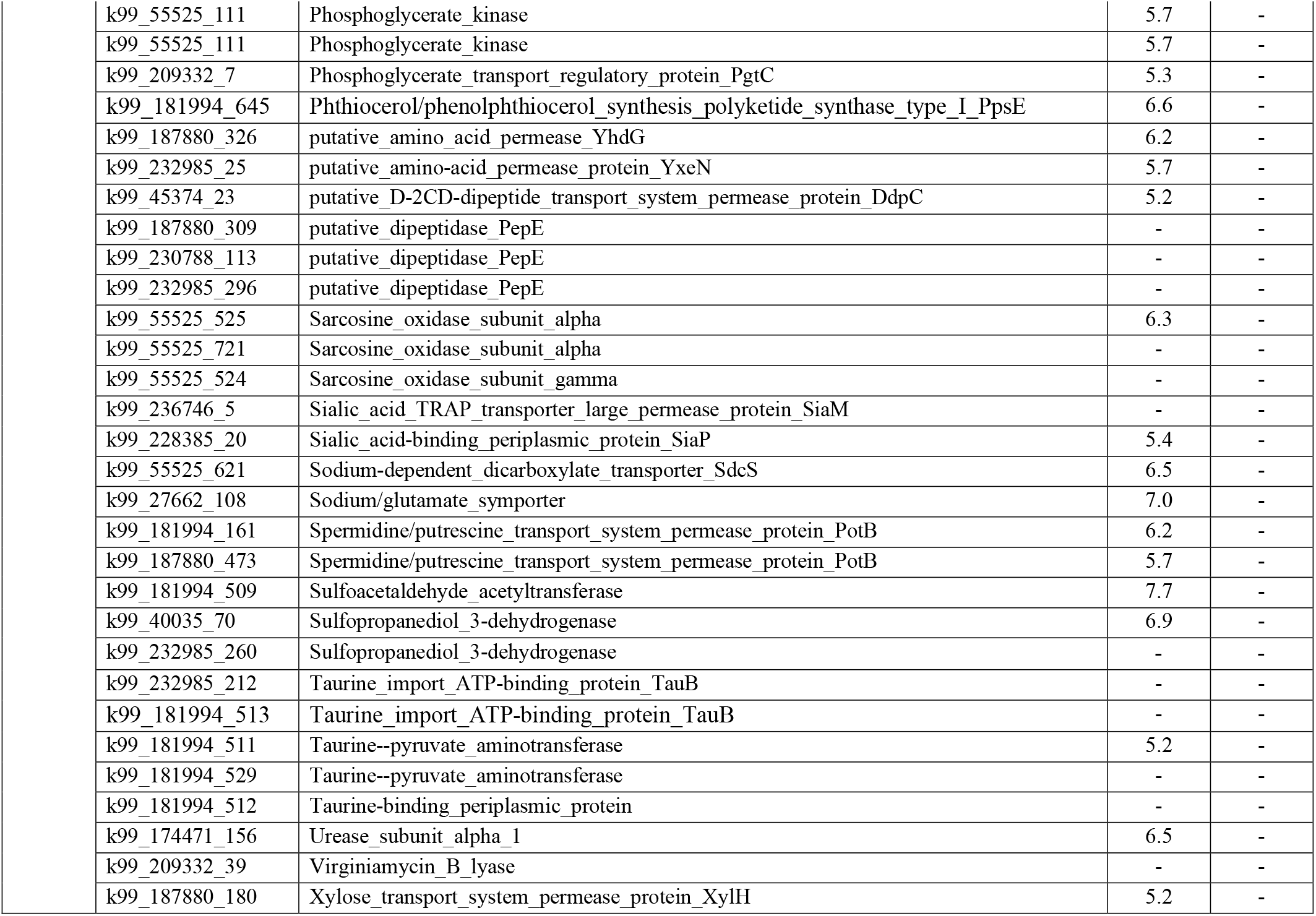
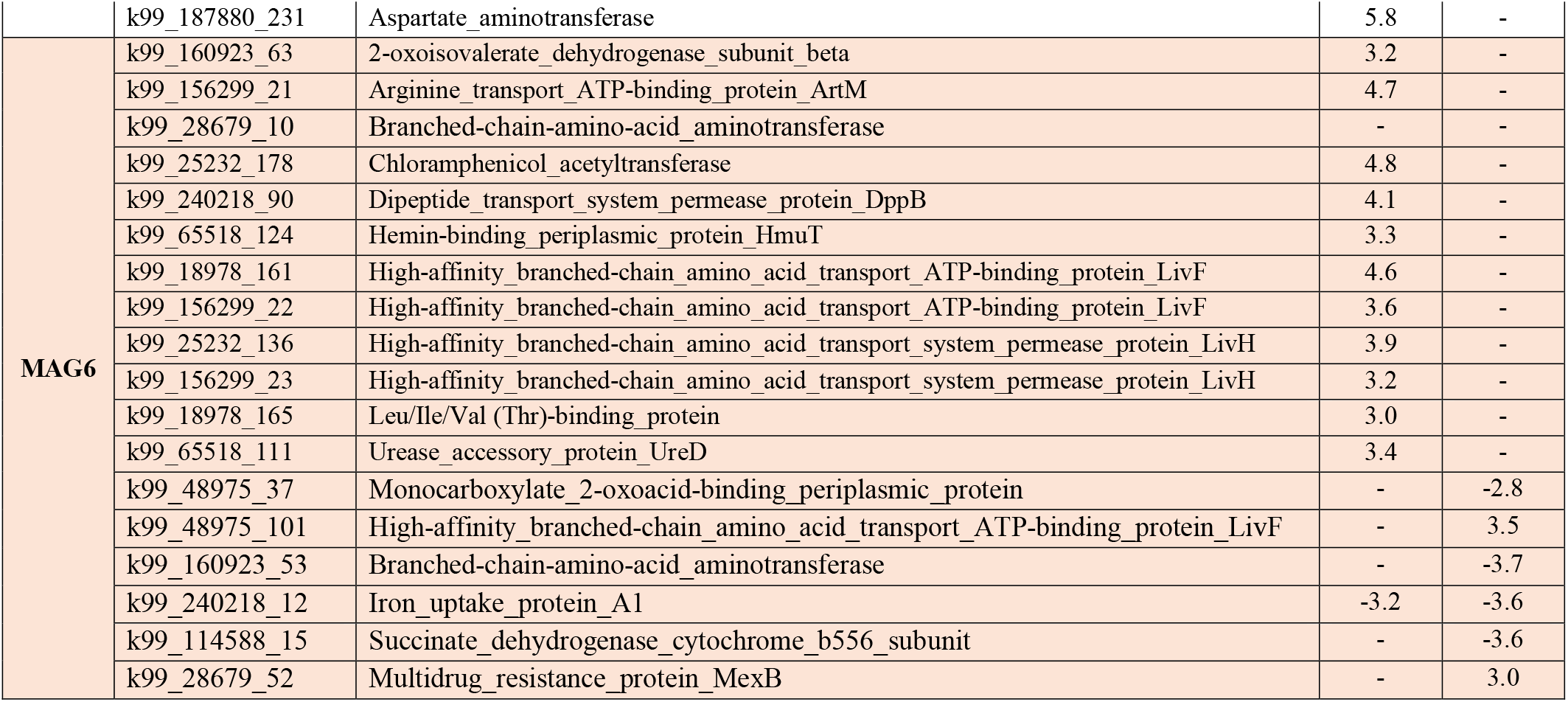
List of selected expressed genes by roseobacter metagenomically-assembled genomes MAG3, MAG5 and MAG6, including genes depicted in Fig. 3. Genes with a false discovery rate (FDR) adjusted *p*-value < 0.1 were considered to be differentially expressed. The values correspond to the log_2_-fold change at the two timepoints in response to reseeding relative to consortium control. Blank cells indicate no differential expression.

| MAG | Gene ID | Annotation | Log<sub>2</sub>-Fold Change |  |
| --- | --- | --- | --- | --- |
|  |  |  | 0.5 hr | 24 hr |
| MAG3 | k99_165394_20 | 2-oxoisovalerate_dehydrogenase_subunit_alpha | - | - |
|  | k99_165394_19 | 2-oxoisovalerate_dehydrogenase_subunit_beta | - | - |
|  | k99_56693_167 | 2-oxoisovalerate_dehydrogenase_subunit_beta | - | - |
|  | k99_258460_5 | Alpha-ketoglutarate-dependent_taurine_dioxygenase | - | - |
|  | k99_157987_224 | Argininosuccinate_lyase | 5.2 | 5.4 |
|  | k99_165394_71 | Bicyclomycin_resistance_protein | 5 | - |
|  | k99_152108_202 | Biofilm_growth-associated_repressor | 6.2 | - |
|  | k99_111971_14 | Branched-chain-amino-acid_aminotransferase | 5.6 | - |
|  | k99_152108_131 | C4-dicarboxylate_TRAP_transporter_large_permease_protein_DctM | 6.6 | - |
|  | k99_87748_267 | C4-dicarboxylate_TRAP_transporter_large_permease_protein_DctM | 6.3 | - |
|  | k99_115381_320 | C4-dicarboxylate_TRAP_transporter_large_permease_protein_DctM | 6.0 | - |
|  | k99_226572_386 | C4-dicarboxylate_TRAP_transporter_large_permease_protein_DctM | 5.6 | - |
|  | k99_165394_264 | Chloramphenicol_acetyltransferase | 5.2 | - |
|  | k99_87748_232 | Creatinase | 6.6 | - |
|  | k99_87748_404 | Dimethylsulfoniopropionate_lyase_DddQ | 6.6 | - |
|  | k99_226572_438 | Flagellar_biosynthetic_protein_FlhB | 6.6 | - |
|  | k99_256486_107 | Glutamate/aspartate_import_permease_protein_GltK | 6.2 | - |
|  | k99_115381_744 | Glutamine_transport_ATP-binding_protein_GlnQ | 5.4 | - |
|  | k99_226572_256 | Heme-binding_protein_A | 6.6 | - |
|  | k99_226572_290 | High-affinity_branched-chain_amino_acid_transport_ATP-binding_protein_LivF | 5.6 | - |
|  | k99_226572_17 | High-affinity_branched-chain_amino_acid_transport_system_permease_protein_LivH | - | - |
|  | k99_226572_237 | High-affinity_branched-chain_amino_acid_transport_system_permease_protein_LivH | 6.1 | - |
|  | k99_152108_173 | High-affinity_branched-chain_amino_acid_transport_system_permease_protein_LivH | 6.0 | - |
|  | k99_94583_24 | High-affinity_branched-chain_amino_acid_transport_system_permease_protein_LivH | 5.7 | - |
|  | k99_166876_14 | Homogentisate_1,2C2-dioxygenase | 6.0 | 6.4 |
|  | k99_166876_81 | Homoserine/homoserine_lactone_efflux_protein | 5.1 | - |
|  | k99_157987_68 | Leucine-responsive_regulatory_protein | 6.0 | - |
|  | k99_157987_182 | Multidrug_export_protein_AcrF | 7.5 | - |
|  | k99_152108_327 | Multidrug_resistance_protein_MdtA | 6.2 | - |
|  | k99_152108_446 | Multidrug_resistance_protein_MdtK | 7.6 | - |
|  | k99_115381_25 | Periplasmic_dipeptide_transport_protein | 6.0 | - |
|  | k99_87748_346 | Phosphate_acetyltransferase | - | - |
|  | k99_157987_27 | Putative_aliphatic_sulfonates_transport_permease_protein_SsuC | 6.2 | - |
|  | k99_115381_555 | putative_dipeptidase_PepE | 6.6 | - |
|  | k99_165394_144 | Putative_multidrug_export_ATP-binding/permease_protein | 6.7 | - |
|  | k99_258460_64 | Putrescine_transport_system_permease_protein_PotH | 5.4 | - |
|  | k99_115381_678 | Putrescine-binding_periplasmic_protein | 5.2 | - |
|  | k99_226572_141 | Sarcosine_oxidase_subunit_alpha | 6.9 | - |
|  | k99_115381_515 | Sarcosine_oxidase_subunit_alpha | 6.8 | - |
|  | k99_115381_401 | Sarcosine_oxidase_subunit_alpha | 6.6 | - |
|  | k99_19927_12 | Sarcosine_oxidase_subunit_alpha | 5.5 | - |
|  | k99_115381_517 | Sarcosine_oxidase_subunit_beta | 6.0 | - |
|  | k99_115381_664 | Sialic_acid_TRAP_transporter_permease_protein_SiaT | 6.3 | - |
|  | k99_94583_171 | Sialic_acid_TRAP_transporter_permease_protein_SiaT | 5.1 | - |
|  | k99_115381_666 | Sialic_acid-binding_periplasmic_protein_SiaP | 6.1 | - |
|  | k99_56693_271 | Sorbitol_dehydrogenase | - | - |
|  | k99_226572_418 | Spermidine/putrescine_import_ATP-binding_protein_PotA | - | - |
|  | k99_115381_675 | Spermidine/putrescine_import_ATP-binding_protein_PotA | 5.8 | - |
|  | k99_226572_460 | Spermidine/putrescine-binding_periplasmic_protein | 5.5 | - |
|  | k99_87748_422 | Sulfoacetaldehyde_acetyltransferase | 6.1 | - |
|  | k99_94583_81 | Sulfoacetaldehyde_acetyltransferase | 5.2 | - |
|  | k99_152108_210 | Sulfopropanediol_3-dehydrogenase | 6.8 | - |
|  | k99_87748_269 | Sulfopropanediol_3-dehydrogenase | 5.1 | - |
|  | k99_157987_26 | Taurine_import_ATP-binding_protein_TauB | - | - |
|  | k99_157987_24 | Taurine--pyruvate_aminotransferase | - | - |
|  | k99_87748_391 | Taurine--pyruvate_aminotransferase | 6.9 | - |
|  | k99_157987_25 | Taurine-binding_periplasmic_protein | - | - |
|  | k99_94583_9 | Urease_subunit_alpha_1 | 5.9 | 8.4 |
|  | k99_247855_81 | Xaa-Pro_dipeptidase | 6.8 | - |
| MAG5 | k99_236746_15 | 2-oxoisovalerate_dehydrogenase_subunit_alpha | - |  |
|  | k99_236746_16 | 2-oxoisovalerate_dehydrogenase_subunit_beta | - | - |
|  | k99_55525_106 | 2-oxoisovalerate_dehydrogenase_subunit_beta | - | - |
|  | k99_55525_477 | 4-hydroxyphenylpyruvate_dioxygenase | - | - |
|  | k99_55525_659 | 4-hydroxyphenylpyruvate_dioxygenase | - | - |
|  | k99_181994_636 | 6"-hydroxyparomomycin_C_oxidase | 6.2 | - |
|  | k99_45374_273 | Acetylornithine_deacetylase | 7.3 | - |
|  | k99_45374_228 | Aclacinomycin_methylesterase_RdmC | - | - |
|  | k99_187880_412 | AI-2_transport_protein_TqsA | - | - |
|  | k99_232985_80 | Argininosuccinate_lyase | 6.7 | - |
|  | k99_181994_54 | Autoinducer_2_sensor_kinase/phosphatase_LuxQ | 6.1 | - |
|  | k99_27662_156 | Autoinducer_2_sensor_kinase/phosphatase_LuxQ | 5.3 | - |
|  | k99_55525_216 | Bicyclomycin_resistance_protein | 5.5 | - |
|  | k99_187880_435 | Bicyclomycin_resistance_protein | - | - |
|  | k99_181994_278 | Branched-chain-amino-acid_aminotransferase | 5.9 | - |
|  | k99_55525_568 | C4-dicarboxylate_transport_sensor_protein_DctB | 6.9 | - |
|  | k99_209332_6 | C4-dicarboxylate_transport_sensor_protein_DctB | 5.7 | - |
|  | k99_55525_233 | C4-dicarboxylate_transport_sensor_protein_DctB | 5.4 | - |
|  | k99_230788_7 | C4-dicarboxylate_transport_transcriptional_regulatory_protein_DctD | 5.2 | - |
|  | k99_45374_16 | C4-dicarboxylate_TRAP_transporter_large_permease_protein_DctM | 6.8 | - |
|  | k99_46830_4 | Cation/acetate_symporter_ActP | 4.4 | - |
|  | k99_236746_25 | Chemotaxis_protein_CheW | 5.4 | - |
|  | k99_181994_493 | Creatinase | 8.3 | - |
|  | k99_174471_78 | Cypemycin_N-terminal_methyltransferase | - | - |
|  | k99_40035_14 | Dipeptidyl-peptidase_5 | 5.3 | - |

|  |  |  |  |
| --- | --- | --- | --- |
| k99_181994_127 | Flagellar_biosynthesis_protein_FlhA | 7.6 | - |
| k99_181994_129 | Flagellar_biosynthetic_protein_FlhB | 6.9 | - |
| k99_181994_55 | Flagellar_P-ring_protein | - | - |
| k99_181994_140 | Flagellum-specific_ATP_synthase | 5.3 | - |
| k99_181994_413 | Glutamine_transport_ATP-binding_protein_GlnQ | 6.1 | - |
| k99_232985_237 | Glutamine_transport_ATP-binding_protein_GlnQ | 5.5 | - |
| k99_181994_174 | Glutamine-binding_periplasmic_protein | 5.3 | - |
| k99_232985_278 | Hemin_transport_system_permease_protein_HmuU | 5.4 | - |
| k99_174471_8 | Hemin_transport_system_permease_protein_HmuU | - | - |
| k99_46830_30 | Hemin_transport_system_permease_protein_HmuU | - | - |
| k99_45374_386 | High-affinity_branched-chain_amino_acid_transport_ATP-binding_protein_LivF | 6.1 | - |
| k99_181994_33 | High-affinity_branched-chain_amino_acid_transport_system_permease_protein_LivH | 5.8 | - |
| k99_45374_387 | High-affinity_branched-chain_amino_acid_transport_system_permease_protein_LivH | 5.2 | - |
| k99_232985_19 | High-affinity_branched-chain_amino_acid_transport_system_permease_protein_LivH | 5.2 | - |
| k99_45374_373 | Homogentisate_1,2C2-dioxygenase | 6.2 | - |
| k99_45374_384 | Leucine-2C_isoleucine-2C_valine-2C_threonine-2C_and_alanine-binding_protein | 5.2 | - |
| k99_181994_28 | Leucine-2C_isoleucine-2C_valine-2C_threonine-2C_and_alanine-binding_protein | - | - |
| k99_187880_15 | Low-molecular_weight_cobalt-containing_nitrile_hydratase_subunit_alpha | 5.2 | - |
| k99_45374_378 | Metallo-beta-lactamase_type_2 | 4.9 | - |
| k99_181994_520 | Methyl-accepting_chemotaxis_protein_II | 6.0 | - |
| k99_181994_779 | Methyl-accepting_chemotaxis_protein_III | 5.6 | - |
| k99_55525_831 | Monocarboxylate_2-oxoacid-binding_periplasmic_protein | 4.7 | - |
| k99_55525_603 | Multidrug_resistance_protein_MdtK | 5.5 | - |
| k99_55525_274 | N-acyl_homoserine_lactonase | 6.8 | - |
| k99_228385_23 | p-hydroxyphenylacetate_3-hydroxylase-2C_reductase_component | - | - |
| k99_181994_654 | Periplasmic_dipeptide_transport_protein | 6.2 | - |
| k99_27662_31 | Periplasmic_dipeptide_transport_protein | 6.0 | - |
| k99_174471_139 | Periplasmic_dipeptide_transport_protein | - | - |
| k99_181994_507 | Phosphate_acetyltransferase | - | - |
| k99_55525_55 | Phosphate_acetyltransferase | - | - |
| k99_55525_111 | Phosphoglycerate_kinase | 5.7 | - |
| k99_55525_111 | Phosphoglycerate_kinase | 5.7 | - |
| k99_209332_7 | Phosphoglycerate_transport_regulatory_protein_PgtC | 5.3 | - |
| k99_181994_645 | Phthiocerol/phenolphthiocerol_synthesis_polyketide_synthase_type_I_PpsE | 6.6 | - |
| k99_187880_326 | putative_amino_acid_permease_YhdG | 6.2 | - |
| k99_232985_25 | putative_amino-acid_permease_protein_YxeN | 5.7 | - |
| k99_45374_23 | putative_D-2CD-dipeptide_transport_system_permease_protein_DdpC | 5.2 | - |
| k99_187880_309 | putative_dipeptidase_PepE | - | - |
| k99_230788_113 | putative_dipeptidase_PepE | - | - |
| k99_232985_296 | putative_dipeptidase_PepE | - | - |
| k99_55525_525 | Sarcosine_oxidase_subunit_alpha | 6.3 | - |
| k99_55525_721 | Sarcosine_oxidase_subunit_alpha | - | - |
| k99_55525_524 | Sarcosine_oxidase_subunit_gamma | - | - |
| k99_236746_5 | Sialic_acid_TRAP_transporter_large_permease_protein_SiaM | - | - |
| k99_228385_20 | Sialic_acid-binding_periplasmic_protein_SiaP | 5.4 | - |
| k99_55525_621 | Sodium-dependent_dicarboxylate_transporter_SdcS | 6.5 | - |
| k99_27662_108 | Sodium/glutamate_symporter | 7.0 | - |
| k99_181994_161 | Spermidine/putrescine_transport_system_permease_protein_PotB | 6.2 | - |
| k99_187880_473 | Spermidine/putrescine_transport_system_permease_protein_PotB | 5.7 | - |
| k99_181994_509 | Sulfoacetaldehyde_acetyltransferase | 7.7 | - |
| k99_40035_70 | Sulfopropanediol_3-dehydrogenase | 6.9 | - |
| k99_232985_260 | Sulfopropanediol_3-dehydrogenase | - | - |
| k99_232985_212 | Taurine_import_ATP-binding_protein_TauB | - | - |
| k99_181994_513 | Taurine_import_ATP-binding_protein_TauB | - | - |
| k99_181994_511 | Taurine--pyruvate_aminotransferase | 5.2 | - |
| k99_181994_529 | Taurine--pyruvate_aminotransferase | - | - |
| k99_181994_512 | Taurine-binding_periplasmic_protein | - | - |
| k99_174471_156 | Urease_subunit_alpha_1 | 6.5 | - |
| k99_209332_39 | Virginiamycin_B_lyase | - | - |
| k99_187880_180 | Xylose_transport_system_permease_protein_XylH | 5.2 | - |
|  | k99_187880_231 | Aspartate_aminotransferase | 5.8 | - |
| MAG6 | k99_160923_63 | 2-oxoisovalerate_dehydrogenase_subunit_beta | 3.2 | - |
|  | k99_156299_21 | Arginine_transport_ATP-binding_protein_ArtM | 4.7 | - |
|  | k99_28679_10 | Branched-chain-amino-acid_aminotransferase | - | - |
|  | k99_25232_178 | Chloramphenicol_acetyltransferase | 4.8 | - |
|  | k99_240218_90 | Dipeptide_transport_system_permease_protein_DppB | 4.1 | - |
|  | k99_65518_124 | Hemin-binding_periplasmic_protein_HmuT | 3.3 | - |
|  | k99_18978_161 | High-affinity_branched-chain_amino_acid_transport_ATP-binding_protein_LivF | 4.6 | - |
|  | k99_156299_22 | High-affinity_branched-chain_amino_acid_transport_ATP-binding_protein_LivF | 3.6 | - |
|  | k99_25232_136 | High-affinity_branched-chain_amino_acid_transport_system_permease_protein_LivH | 3.9 | - |
|  | k99_156299_23 | High-affinity_branched-chain_amino_acid_transport_system_permease_protein_LivH | 3.2 | - |
|  | k99_18978_165 | Leu/Ile/Val (Thr)-binding_protein | 3.0 | - |
|  | k99_65518_111 | Urease_accessory_protein_UreD | 3.4 | - |
|  | k99_48975_37 | Monocarboxylate_2-oxoacid-binding_periplasmic_protein | - | -2.8 |
|  | k99_48975_101 | High-affinity_branched-chain_amino_acid_transport_ATP-binding_protein_LivF | - | 3.5 |
|  | k99_160923_53 | Branched-chain-amino-acid_aminotransferase | - | -3.7 |
|  | k99_240218_12 | Iron_uptake_protein_A1 | -3.2 | -3.6 |
|  | k99_114588_15 | Succinate_dehydrogenase_cytochrome_b556_subunit | - | -3.6 |
|  | k99_28679_52 | Multidrug_resistance_protein_MexB | - | 3.0 |

**Table S6.**
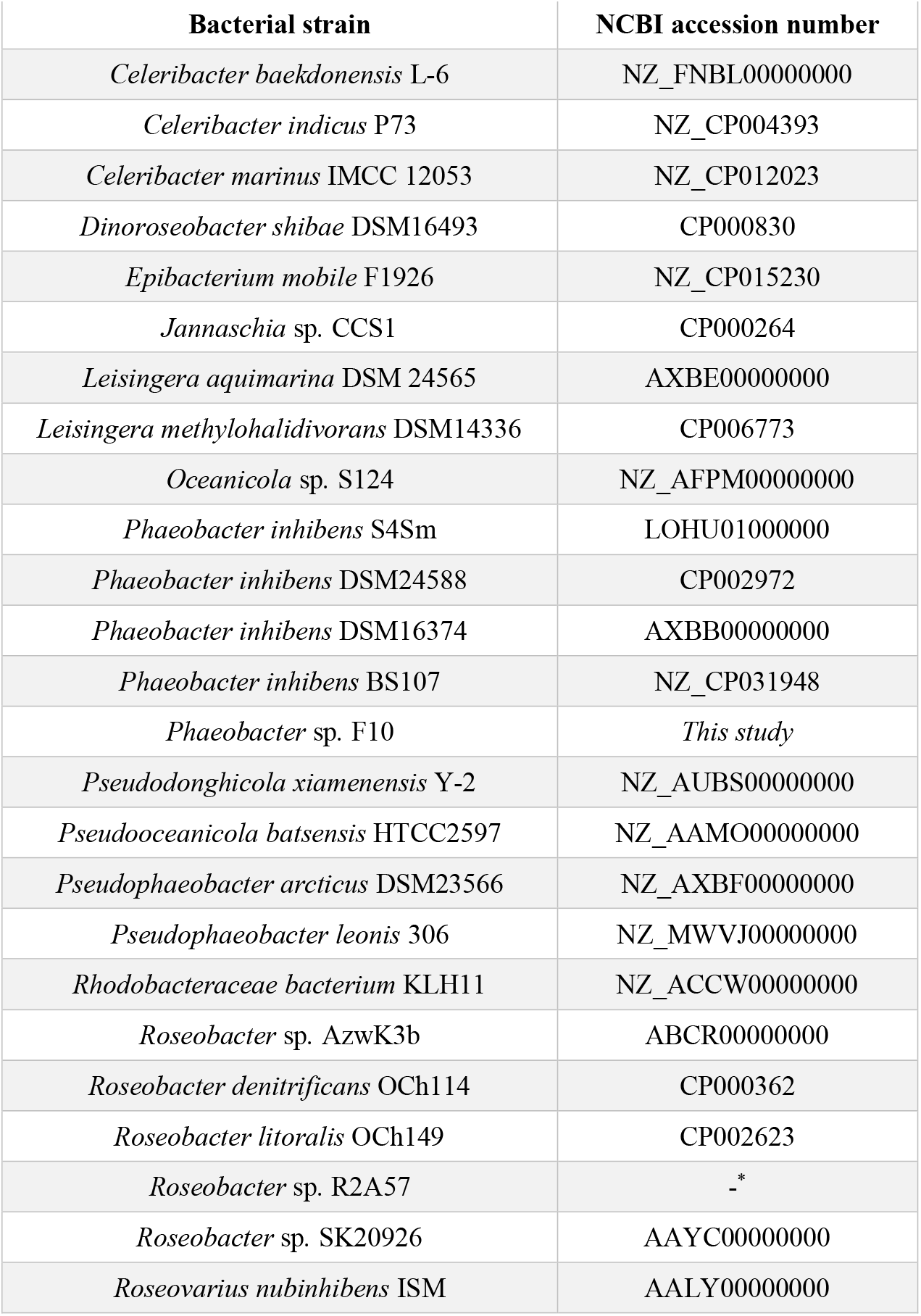
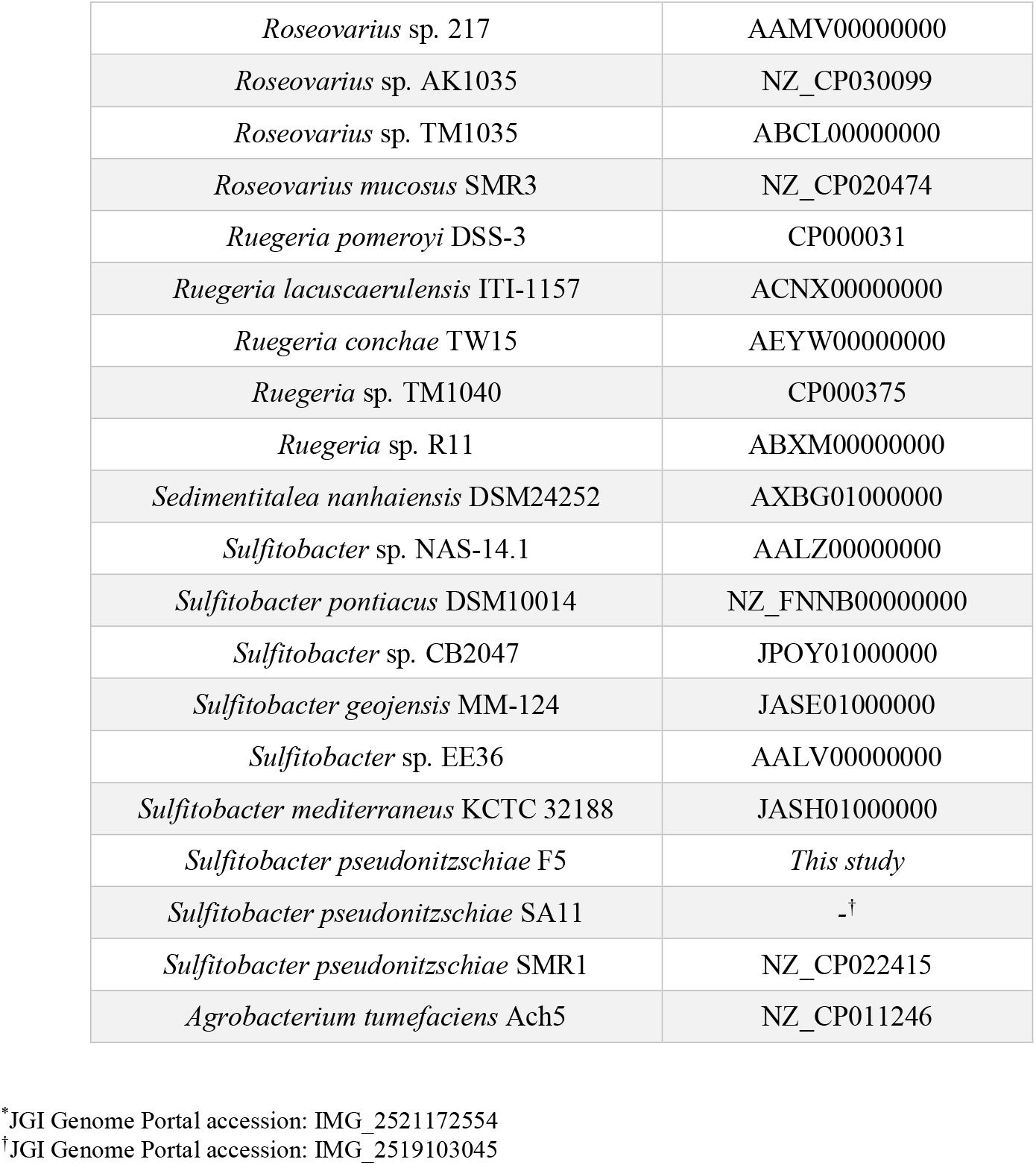
List of accession numbers for roseobacter genomes used in the phylogenomic analysis of the roseobacter MAGs and laboratory cultured consortium strains (Fig. S3). The *Agrobacterium tumefaciens* Ach5 genome was used as an outgroup.

**Table S7.**
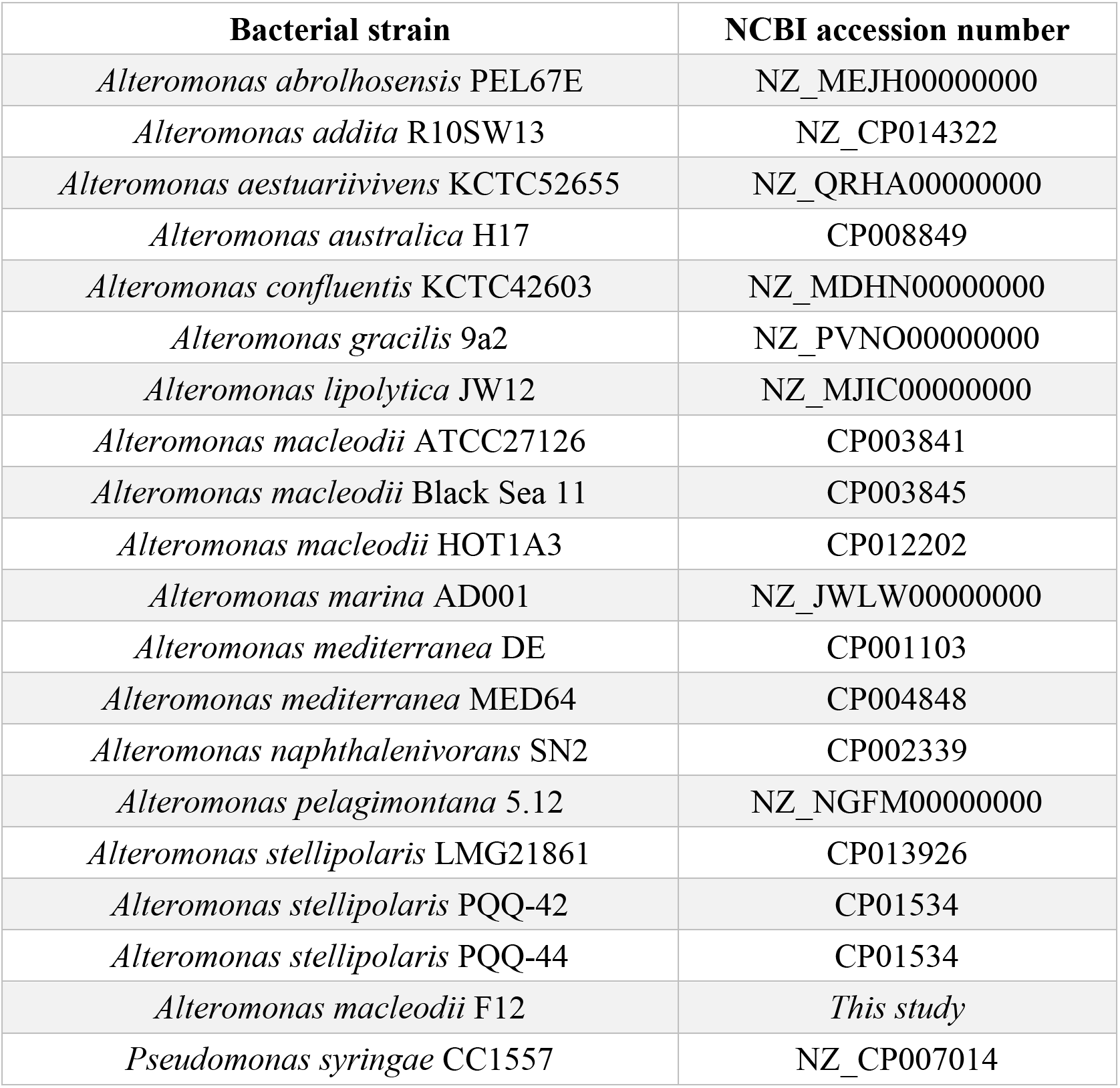
List of accession numbers for Alteromonadaceae genomes used in the phylogenomic analysis of the Alteromonadaceae MAGs and a laboratory cultured consortium strain (Fig. S4). *Pseudomonas syringae* CC1557 genome was used as an outgroup.

**Table S8.** High-throughput sequencing information and read counts for the *A. glacialis* A3 genome (A3Ax), *A. glacialis* A3 RNA-seq samples (D1-D12), bacterial consortium metagenome (A3Bact), and bacterial RNA-seq samples (B1-B9).

| Sample name | Description | Number of reads (raw) | Number of quality trimmed reads | Retained (%) | Lost (%) |
| --- | --- | --- | --- | --- | --- |
| A3Ax | Diatom genome | 165811606 | 153290706 | 92.4 | 7.6 |
| D1 | Diatom only at 0.5 hours - rep 1 | 55597046 | 45927566 | 82.6 | 17.4 |
| D2 | Diatom only at 0.5 hours - rep 2 | 32696296 | 25491292 | 78.0 | 22.0 |
| D3 | Diatom only at 0.5 hours - rep 3 | 27352918 | 21994556 | 80.4 | 19.6 |
| D4 | Diatom only at 24 hours - rep 1 | 41701950 | 33661898 | 80.7 | 19.3 |
| D5 | Diatom only at 24 hours - rep 2 | 31989840 | 24022442 | 75.1 | 24.9 |
| D6 | Diatom only at 24 hours - rep 3 | 27881510 | 23237124 | 83.3 | 16.7 |
| D7 | Reseeded diatom at 0.5 hours - rep 1 | 26184068 | 20373958 | 77.8 | 22.2 |
| D8 | Reseeded diatom at 0.5 hours - rep 2 | 32222538 | 24054500 | 74.7 | 25.3 |
| D9 | Reseeded diatom at 0.5 hours - rep 3 | 37628278 | 29615596 | 78.7 | 21.3 |
| D10 | Reseeded diatom at 24 hours - rep 1 | 29849476 | 23719096 | 79.5 | 20.5 |
| D11 | Reseeded diatom at 24 hours - rep 2 | 32760398 | 26860754 | 82.0 | 18.0 |
| D12 | Reseeded diatom at 24 hours - rep 3 | 32198030 | 25647272 | 79.7 | 20.3 |
| A3Bact | Bacterial consortium metagenome | 905113782 | 828297028 | 91.5 | 8.5 |
| B1 | Consortium only at 0.5 hours - rep 1 | 204707556 | 135964890 | 66.4 | 33.6 |
| B2 | Consortium only at 0.5 hours - rep 2 | 123255494 | 75388906 | 61.2 | 38.8 |
| B3 | Consortium only at 0.5 hours - rep 3 | 106506474 | 66325910 | 62.3 | 37.7 |
| B4 | Reseeded consortium at 0.5 hours - rep 1 | 129606370 | 56559286 | 43.6 | 56.4 |
| B5 | Reseeded consortium at 0.5 hours - rep 2 | 103087344 | 79271378 | 76.9 | 23.1 |
| B6 | Reseeded consortium at 0.5 hours - rep 3 | 50236330 | 32005540 | 63.7 | 36.3 |
| B8 | Reseeded consortium at 24 hours - rep 1 | 41267162 | 12780816 | 31.0 | 69.0 |
| B9 | Reseeded consortium at 24 hours - rep 2 | 205670530 | 146660984 | 71.3 | 28.7 |

**Dataset S1.** (Excel CSV format) List of retention times (RT) and mass-to-charge values (m/z) for axenic and reseeded samples at four timepoints (0.5, 4, 24, and 48 hours) analyzed on an ultrahigh-performance liquid chromatography quadrupole time-of-flight mass spectrometer (UHPLC-QToF-MS).

**Dataset S2.** (Excel CSV format) List of expected masses (m/z) (ExpMass), peak intensities, theoretical masses (ThMass) and chemical formulae for the DOM composition in axenic samples using a Fourier-transform ion cyclotron resonance mass spectrometry (FT-ICR-MS).

**Dataset S3.** (Excel CSV format) List of expected masses (m/z) (ExpMass), peak intensities, theoretical masses (ThMass) and chemical formulae for the DOM composition in reseeded samples using a Fourier-transform ion cyclotron resonance mass spectrometry (FT-ICR-MS).

